# A new drug candidates for glycogen storage disorders enhances glycogen catabolism: Lessons from Adult Polyglucosan Body Disease models

**DOI:** 10.1101/2021.03.18.436069

**Authors:** Hilla Vaknin, Kumudesh Mishra, Jeevitha D’Souza, Monzer Marisat, Uri Sprecher, Shane Wald-Altman, Anna Dukhovny, Yuval Raviv, Benny Da’adoosh, Hamutal Engel, Sandrine Benhamron, Keren Nitzan, Anna Permyakova, Hanna Rosenmann, Alexander Lossos, Joseph Tam, Berge A. Minassian, Or Kakhlon, Miguel Weil

**Author notes:** Equal contribution (Hilla Vaknin, Kumudesh Mishra, Jeevitha D’Souza).

## Abstract

This work employs Adult Polyglucosan Body Disease (APBD) models to explore the efficacy and mechanism of action of 144DG11, a new polyglucosan-reducing lead compound discovered by a high-throughput screen (HTS). APBD is an adult onset glycogen storage disorder (GSD) manifesting as a debilitating progressive axonopathic leukodystrophy. APBD is caused by glycogen branching enzyme (GBE) deficiency leading to poorly branched and insoluble glycogen inclusions, which precipitate as neuropathogenic polyglucosans (PG). 144DG11 led to prolonged survival and improved motor parameters in a GBE knockin (Gbe^ys/ys^) APBD mouse model. Histopathologically, 144DG11 reduced PG and glycogen levels in brain, liver, heart, and peripheral nerve. Indirect calorimetry experiments revealed that 144DG11 increases carbohydrate burn at the expense of fat burn, suggesting metabolic mobilization of pathogenic PG. These results were also reflected at the cellular level by increased glycolytic, mitochondrial and total ATP production. Mechanistically, we show that the molecular target of 144DG11 is the lysosomal membrane protein LAMP1, whose interaction with the compound, similar to LAMP1 knockdown, enhanced autolysosomal degradation of glycogen and lysosomal acidification. Enhanced mitochondrial activity and lysosomal modifications were also the most pronounced effects of 144DG11 in APBD patient fibroblasts as discovered by image-based multiparametric phenotyping analysis and corroborated by proteomics. In summary, this work presents a broad mechanistic and target-based characterization of 144DG11 in in vivo and cell models of the prototypical GSD APBD. This investigation warrants development of 144DG11 into a safe and efficacious GSD therapy.

**One Sentence Summary:** A new compound, demonstrated to ameliorate APBD in vivo and ex vivo by autophagic catabolism of glycogen, may potentially become a universal drug for glycogen storage disorders.

## Introduction

Adult Polyglucosan Body Disease (APBD), is a glycogen storage disorder (GSD) which manifests as a debilitating and fatal progressive axonopathic leukodystrophy from the age of 45-50 (*1, 2*). APBD is further characterized by peripheral neuropathy, dysautonomia, urinary incontinence and occasionally dementia, all being important diagnostic criteria for this commonly misdiagnosed (*3, 4*) and widely heterogeneous disease. APBD is caused by glycogen branching enzyme (GBE) deficiency leading to poorly branched and therefore insoluble glycogen (polyglucosans, PG), which precipitate, aggregate and accumulate into PG bodies (PB). Being out of solution and aggregated, PB cannot be digested by glycogen phosphorylase. The amassing aggregates lead to liver failure and death in childhood (Andersen’s disease; GSD type IV). Milder mutations of GBE, such as p.Y329S in APBD, lead to smaller PB, which do not disturb hepatocytes and most other cell types, merely accumulating in the sides of cells. In neurons and astrocytes, however, over time PB plug the tight confines of axons and processes and lead to APBD.

The majority of APBD patients are Ashkenazi Jewish bearing the p.Y329S mutation in the *Gbe1* gene (similar to the knockin APBD modeling mice we use here (*5*)). More than 200 confirmed APBD cases worldwide have been reported to date. However, due to common misdiagnosis, this is probably an underestimation. In addition, APBD’s carrier frequency is relatively high (1/58 deduced from testing 2,776 individuals self-reported to be 100% Ashkenazi Jewish (*4, 6*), *cf.* 1/27-1/30 carrier frequency in Tay Sachs disease) and therefore, its prevalence is probably underestimated. Moreover, awareness for the disease among physicians and neurologists has only developed in the last decade.

While an effective cure for APBD is urgently needed, APBD is representative of the larger group of GSDs. GSDs are a varied group of 15 incurable diseases with a combined frequency of 1 in 20,000-43,000 (*7*). Ranging from childhood liver disorders such as GSD1, through adolescent myoclonic epilepsies such as Lafora Disease (LD), and adult progressive neurodegenerative disorders such as APBD, all GSDs are currently incurable. A notable exception is Pompe disease, for which acid alpha-glucosidase enzyme replacement therapy (*8, 9*), and recently the more advanced antibody-enzyme fusion therapy (*10, 11*), are promising approaches, albeit with some immunological complications.

Since all GSDs share the etiology of excessive normal or malconstructed glycogen, we believe that APBD represents a prototypical GSD and thus development of pharmaceutical inhibitors of glycogen accumulation is likely to benefit all GSDs (except for GSD0, where glycogen deficiency, rather than surplus, is considered the pathogenic factor (*12*)). Thus, based on the premise that glycogen, or insoluble glycogen in the case of APBD, is a pathogenic factor in GSDs, we have developed an APBD patient cell-based assay to identify small molecule inhibitors of accumulation of insoluble glycogen, or PB (*13*). Applying this assay in a high throughput format, we have screened FDA approved and de novo synthesized compound libraries to discover PB reducing hits, which were further investigated in vitro and in vivo. One of these hits, the FDA approved glycogen synthase inhibitor guaiacol, indeed demonstrated in vivo safety and partial efficacy in the Gbe knockin mouse model of APBD Gbe^ys/ys^ (*14*).

Here we describe the development of a more promising glycogen lowering HTS hit, 144DG11. Compared to guaiacol, whose main therapeutic effect besides polyglucosan lowering was lifespan extension, 144DG11 also improved motor parameters and carbohydrate burn in Gbe^ys/ys^ mice. Furthermore, our extensive cell and in vitro analyses show that 144DG11 targets the lysosomal protein LAMP1 and thus enhances glycophagy and respiratory and glycolytic ATP production, transforming the adversely excessive glycogen into a fuel source. In summary, this study uncovers a new safe and efficacious agent for ameliorating APBD and potentially other GSDs instigated by glycogen over-accumulation.

## Results

### The safe compound 144DG11 improves survival and motor deficiencies in Gbe^ys/ys^ mice

We tested 144DG11 (Fig. 1A) for its capacity to correct the deficient motor phenotypes and short lifespan in the APBD mouse model Gbe^ys/ys^. 144DG11 is one of 19 PG reducing HTS hits previously discovered by us (*13*). It was shown by in silico ADMET (Absorption, Distribution, Metabolism, Excretion, and Toxicity) studies to be safe and pharmacokinetically and pharmacodynamically preferred and is therefore worth further pursuit (Fig. S1). Indeed, low ADMET scoring compounds such as “B” (Fig. S1), were not efficacious and caused adverse effects such as wounds (Fig. S2). Moreover, safety assessment in wild type (wt) mice confirmed that, administered for 3 months at 250 mg/kg in 5% DMSO (the highest dose possible due to solubility and DMSO toxicity issues), 144DG11 did not influence animals’ weight gain over time (Fig. S3). The compound also did not produce any histopathological damage or lesions in brain, liver, skeletal muscle and heart after 3 month exposure (Fig. S4). Following 1h and 24h treatments, mice were also examined for abnormal spontaneous behavior, such as immobility, excessive running, stereotyped movements, and abnormal posture (Irwin tests). 144DG11 did not cause any adverse effect in these Irwin tests (Table S1).

**Fig. 1.**
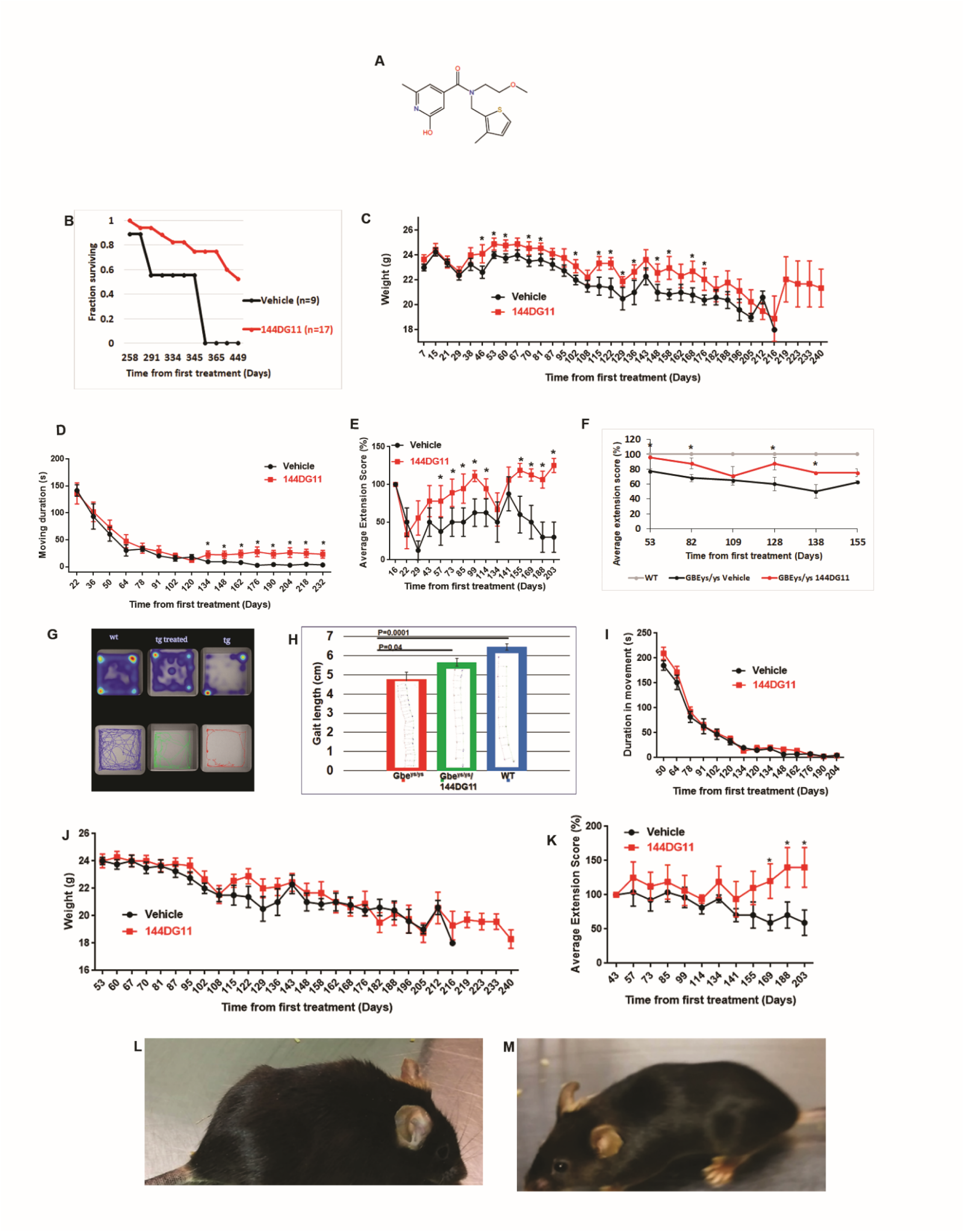
In vivo effects of 144DG11. **(A)** Structural formula of 144DG11. **(B)** Kaplan-Meier survival curve based on 17 animals treated twice a week with 250 mg/kg 144DG11 compared to 9 animals treated with 5% DMSO vehicle. 144DG11 significantly increased survival (log-rank test p-value < 0.000692). **(C)** Weight (in g), **(D)** average duration in movement in an open field, and **(E and F)** extension reflex (the degree to which the hind paws open after holding the animal from the tail) as a function of time after treating wt (n=8) mice with vehicle and Gbe^ys/ys^ mice with vehicle (n=8), or 144DG11 (n=9) as indicated. All animals were treated twice a week with 250 mg/kg from 4 months of age. Two way ANOVA with repeated measures show that throughout the period weight in treatment was higher than vehicle in **(C)** (p<0.05), **(D)** (p<0.1), **(E)** (p<0.05) and **(F)** (p<0.05). Bonferroni’s multiple comparisons test showed no significant difference (at p<0.05) between vehicle and treatment over all time points. Significant difference at specific time points is denoted by *. In **(F)** this difference was significant at all time points except 109 days after first injection. **(G)** Movement Heat Map (*upper panel*, average based on n=9 nine month old females) showing quantification of Open Field performance experiments. Time spent at location is quantified as a heat map (see blue (low) to red (high) scale at bottom right). *Lower panel* shows visual tracking examples from single animals. wt, Untreated wt animals as controls; tg treated, Gbe^ys/ys^ (transgenic) mice treated with 144DG11; tg, APBD mice treated with vehicle. **(H)** Gait analysis (stride length) of n=9 nine months old female mice from each arm (red, Gbe^ys/ys^ mice vehicle treated; green, Gbe^ys/ys^ mice treated with 144DG11; blue, wt, vehicle treated mice). Shown are average (+/- s.d.) stride lengths. A representative ink foot-print trail is also shown for each arm. Statistically significant differences, as tested by One Way ANOVA with Sidak’s post-hoc tests, were demonstrated between 144DG11 and vehicle treated transgenic mice, and between transgenic and wt mice, as indicated by the levels of significance shown. **(I)** Average duration in movement in an open field, **(J)** weight (in g), and **(K)** extension reflex as a function of time after treating Gbe^ys/ys^ mice with vehicle or 144DG11as indicated at 6 months of age (at onset). Two way ANOVA with repeated measures showed that treatment was not different from vehicle over time for **(I)** and **(J)** (p<0.15) and was significant only with p<0.07 for **(K)**. Bonferroni’s multiple comparisons test showed no significant difference (p<0.05) between vehicle and treatment at all time points for **(I)**, **(J)** and **(K)**. 7 month old Gbe^ys/ys^ mice were treated with vehicle **(L)**, or 144DG11 **(M)** for 3 months prior to photography. Note unkempt fur in vehicle treated animal, which was improved by 144DG11. *, Significant difference (p<0.05) for a single time comparison.

Importantly, as Fig. 1B shows, treatment with 144DG11 significantly improved animal survival (log-rank test p-value < 0.000692) compared to vehicle treated animals. Lifespan extension probably mirrors improvement of several parameters related to ability to thrive. The most prominent parameter in that respect is weight. 144DG11 indeed mitigated the decline in animal weight over time caused by the disease (Fig. 1C). We also tested the effect of 144DG11 on various motor parameters. 144DG11 improved open field performance (Fig. 1D) from a relatively advanced stage of disease progression (8 months, 134 days post injection (Fig. 1D)). These improvements were manifested as increased locomotion and an increased tendency to move towards the center (Fig. 1G), perhaps also associated with amelioration of stress and anxiety. The progressive deterioration of Gbe^ys/ys^ mice in open field performance is related to their gait deficiency. Therefore, we tested the effect of 144DG11 on gait at the age of 9 months when gait is severely affected. At that age, 144DG11 indeed improved gait, or increased stride length (Fig. 1H). Our data also show that, of all motor parameters tested, the most pronounced ameliorating effect was on the overall extension reflex (Fig. 1E, F). Overall extension reflex throughout the study period, was significantly improved by 144DG11 (Fig. 1F, p<0.05) as it was at 9 specific time points (asterisks in Fig. 1F). This effect is especially important since its human patient correlate is pyramidal tetraparesis, or upper motor neuron signs, which are one of the main neurological deficiencies in APBD (*2*). Importantly, while open field performance (Fig. 1G), gait (Fig. 1H) and extension reflex (Fig. 1E, F) were significantly improved by 144DG11, they were not restored to wt levels, demonstrating that while efficacious, 144DG11 performance still leaves some room for future improvement.

These effects of 144DG11 on motor parameters were studied by administrating 144DG11 at the age of 4 months, two months prior to disease onset, assuming a preferred prophylactic effect. Such an effect is expected in a neurodegenerative disorder such as APBD in which the already dead neurons cannot be affected by a post-onset treatment. This assumption was validated as for all the parameters improved by 144DG11 - open field (Fig. 1I), weight (Fig. 1J) and overall extension reflex (Fig. 1K) - its ameliorating effect did not take place when administered after disease onset at the age 6 months. Notably, extension reflex, the parameter most affected by 144DG11, was also the only parameter improved by the compound from the advanced stage of the disease at the age of 9 months (Fig. 1K). The overall beneficial effect of 144DG11 can be best appreciated by our supplementary video (Movie S1) and animal photographs which illustrate that treated animals are less kyphotic and better kempt (Figs. 1L & 1M).

### *144DG11* reduces histopathological accumulation of polyglucosans and glycogen in accordance with its biodistribution

As 144DG11 significantly improved motor and survival parameters, we set out to investigate its histopathological effects. This information is important for determining whether the expected mode of action of 144DG11 discovered ex vivo - reduction of polyglucosan levels in fibroblasts (*13*) – also takes place in vivo and if so in which tissues. Brain, heart, muscle, nerve fascicles (peripheral nerves), and liver tissues from 144DG11 and vehicle treated animals were collected following animal sacrificing at age 9.5 months. The same tissues from wt mice were used as controls. Following diastase treatment to digest non-polyglucosan glycogen, leaving behind polyglucosan, sections were stained for polyglucosan with periodic acid-Schiff’s (PAS) reagent, counterstained with hematoxylin and analyzed by light microscopy. The results (Fig. 2A) show a significant reduction in polyglucosan levels in brain, liver, heart, and peripheral nerve, with no apparent effect on muscle polyglucosans. Total glycogen levels, determined biochemically as described in (*15*), were also correspondingly affected (Fig. 2B). These results could possibly explain the improvement observed in motor parameters and in animal thriving (Fig. 1).

**Fig. 2.**
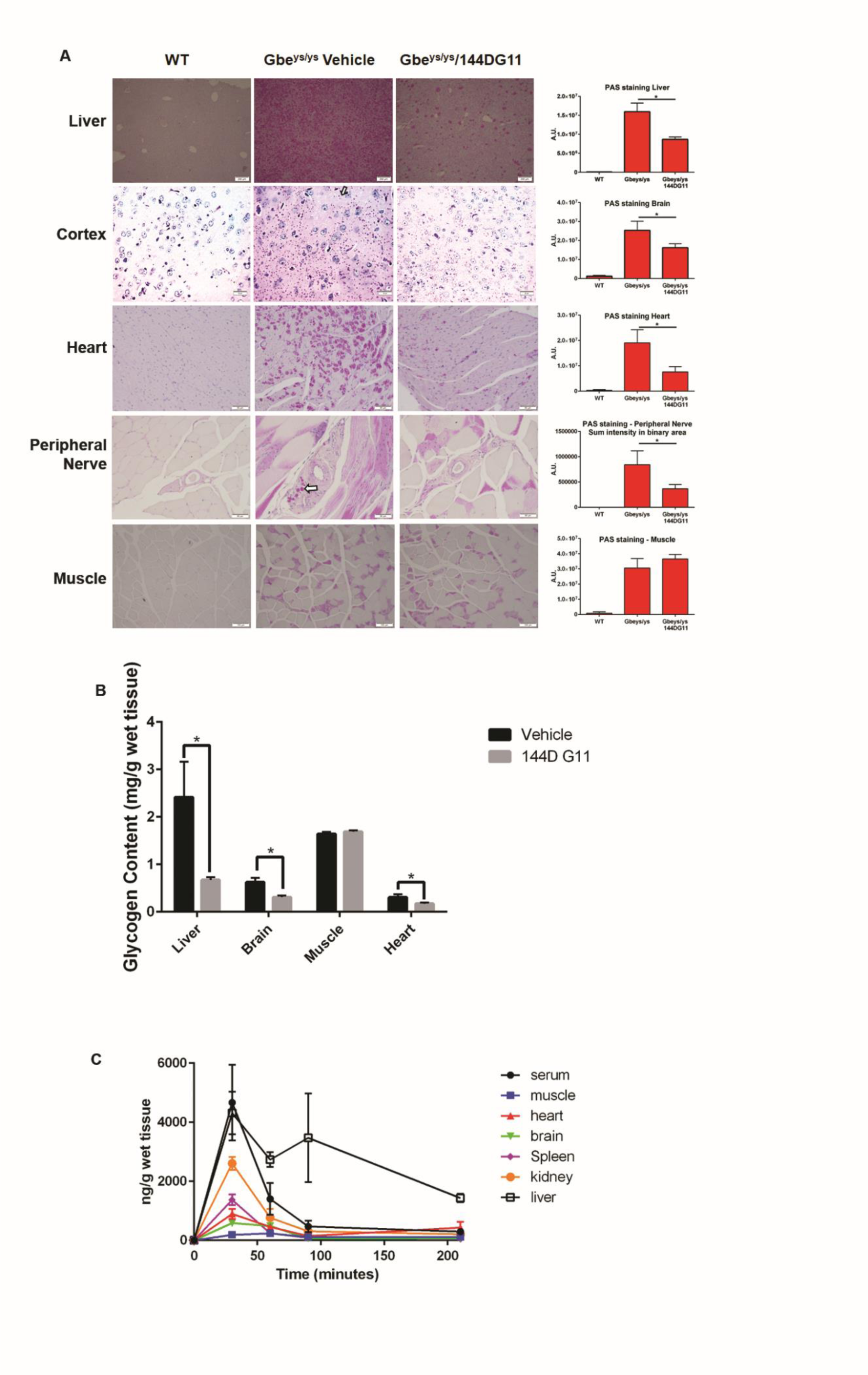
Histopathological effects of 144DG11 and its pharmacokinetics. **(A)** *Left panel,* Mice, treated as indicated above, were sacrificed and the indicated tissues were removed and stained for PG (arrows) with PAS following treatment with diastase. *Right panel,* PAS staining was quantified, as described in (*15*), based on analysis of 4 sections from each tissue in n=3 wt, n=7 Gbe^ys/ys^ vehicle-treated, and n=9 144DG11-treated mice. **(B)** Total glycogen in the respective tissues was quantified as described in (*15*). **(C)** 144DG11 Pharmacokinetics. 9 month old Gbe^ys/ys^ mice were SC-injected with 150 uL of 144DG11 at 250 mg/kg. Mice were sacrificed 30, 60, 90, and 210 min post injection and the indicated tissues were removed, as well as 200 uL of serum drawn. Graph shows 144DG11 levels in the different tissues determined by LC-MS/MS (see Supplementary Materials). Shown are means and SEM of results obtained from n = 3 mice at each time point. Repeated-measures 2-way ANOVA tests show that the pharmacokinetic profile of each tissue is significantly different from that of all other tissues (p < 0.05). **\***, Significant difference (p<0.05) as determined by Student’s t-tests.

Pharmacokinetic analysis is instrumental for explaining the effects of 144DG11 *in situ* regardless of its innate capacity to modify polyglucosans in isolated cells, the reason being that timing of arrival, distribution and stability in the tissue are key determinants of the *in situ* activity of any pharmacological agent. To determine the distribution and kinetic parameters of 144DG11 in different tissues, we treated Gbe^ys/ys^ mice with 250 mg/kg 144DG11 via subcutaneous injection, as in our efficacy experiments. Mice were then sacrificed 0, 30, 60, 90, and 210 min post administration and serum as well as brain, kidney, hind limb skeletal muscle, heart, liver, and spleen tissues were collected, homogenized, extracted, and their 144DG11 levels were analyzed by liquid chromatography tandem mass spectrometry (LC-MS/MS). The results are shown in Fig. 2C. The differential effects of 144DG11 on glycogen and polyglucosan content in the different tissues match its differential distribution and dwell time in each respective tissue. The highest extent of polyglucosan/glycogen reduction was observed in the liver matching the highest dwell time/persistence of 144DG11 observed in the organ (estimated half-life of more than 3 h). The heart and brain demonstrate intermediate levels of 144DG11. However, those levels persist up until 60 minutes post injection, which might account for the 144DG11-mediated reduction in polyglucosan and glycogen content observed in these tissues. The muscle, on the other hand, demonstrated only negligible accumulation of 144DG11, in agreement with lack of effect of the compound on muscle glycogen and polyglucosan content. Based on the sampling times used, time to C_max_ was 30 min for all the tissues studied indicating similar rate of absorption to all these tissues. The highest C_max_ is observed in liver and kidney matching their well-established rapid perfusion. Expectedly, the lowest C_max_ was observed in the skeletal quadriceps muscle, which is known to be a poorly perfused organ.

### *144DG11* enhances carbohydrate metabolism and improves metabolic panel in vivo

The effect of 144DG11 on various metabolic parameters was determined in vivo using indirect calorimetry. Fuel preference at the whole animal level is determined by the respiratory quotient (RQ, the ratio of CO_2_ produced to O_2_ consumed). Lower RQ indicates higher fat burn, while higher RQ indicates higher carbohydrate burn. As our results (Fig. 3A) show, 144DG11 increased RQ to even higher levels than those of the wt animals. The parallel increases, induced by 144DG11, in total energy expenditure (Fig. 3B) and carbohydrate burning at the expense of fat burning (Figs. 3C & 3D) suggest that 144DG11 stimulates glycogen mobilization, which is a therapeutic advantage since Gbe^ys/ys^ mice store glycogen as insoluble and pathogenic polyglucosan. Stimulation of ambulatory activity (Fig. 3E) and of meal size (Fig. 3F) are in line with this observation of stimulation of carbohydrate catabolism in affected animals by 144DG11. Taken together, the increased fuel burning and food intake indicate that 144DG11 can improve metabolic efficiency in the affected animals.

**Fig. 3.**
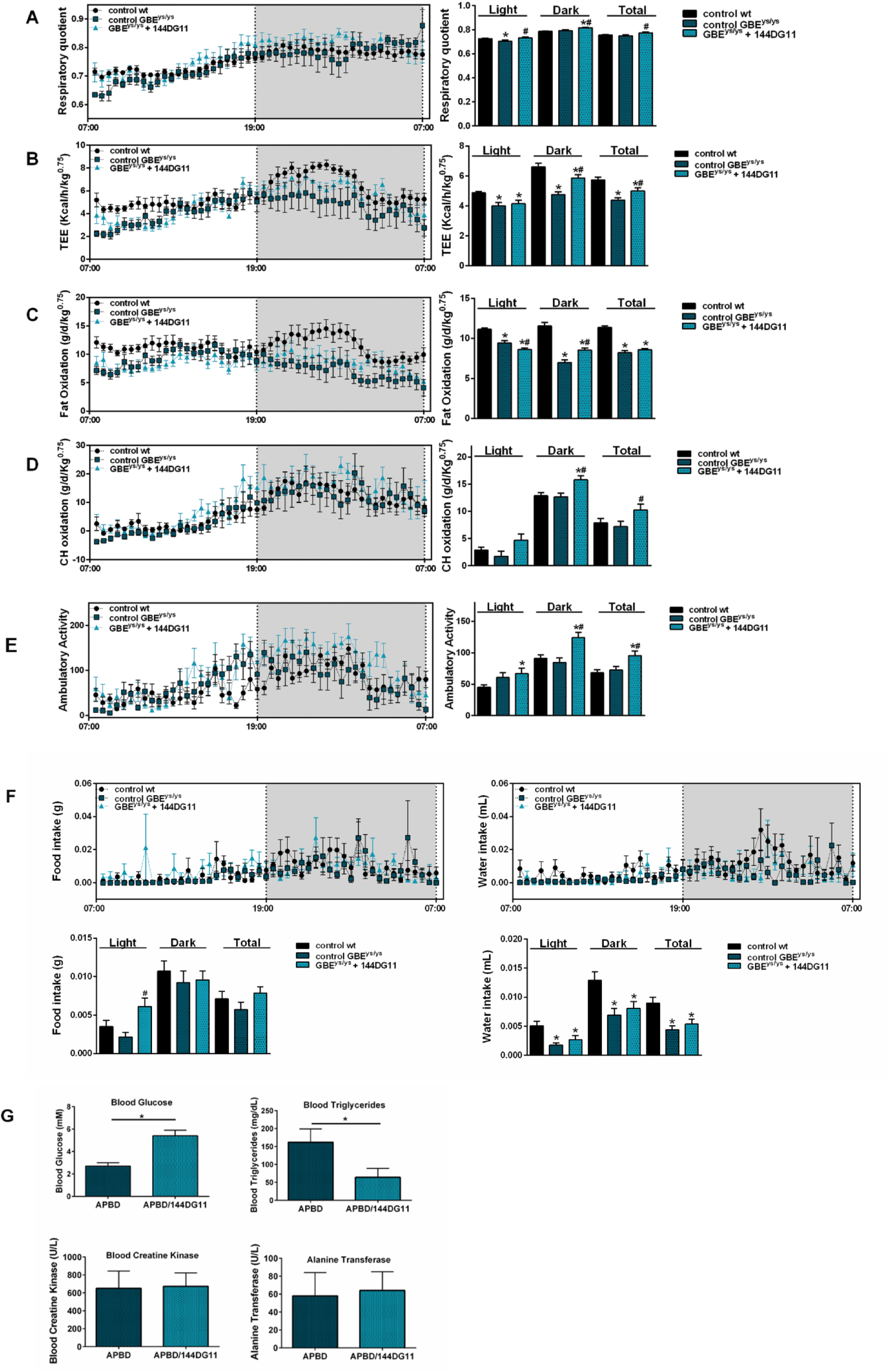
In vivo metabolic profile of mice treated with 144DG11. Mice were monitored over a 24 hr period. Effective mass was calculated by ANCOVA (see Supplementary Materials). Data are mean±SEM from n=11 nine month old mice in the wt control vehicle arm, n=6 nine month old mice in the Gbe^ys/ys^ vehicle arm and n=7 nine month old mice in the Gbe^ys/ys^ 144DG11-treated arm. All injections were from the age of 4 mo. Untreated Gbe^ys/ys^ mice demonstrate *lower* respiratory quotient (in the light) **(A)**, total energy expenditure (TEE) **(B)**, and fat oxidation **(C)** compared to wt controls. 144DG11 treatment increased these parameters (for fat oxidation only in the dark and total time). Carbohydrate oxidation and ambulatory activity, not significantly affected by the diseased state, were increased by 144DG11 even beyond wt control levels **(D and E)** (note, while 144DG11 increased carbohydrate oxidation in the light **(D)**, p was only <0.06). 144DG11 has also reversed the decrease in meal size and water sip volume observed in Gbe^ys/ys^ mice as compared to wt control **(F)**. **(G)** Blood metabolic panel based on n=5, 9.5 month old mice treated as indicated. Blood glucose was increased and blood triglycerides decreased in Gbe^ys/ys^ cells by 144DG11 (p<0.05). *p<0.05 *v* wt controls, #p<0.05 *v* Gbe^ys/ys^ vehicle treated mice. Statistical differences were determined by Student’s t-tests.

We further tested whether 144DG11 is able to correct the hypoglycemia and hyperlipidemia observed in Gbe^ys/ys^ mice (*5*). Such an effect is expected from an agent capable of inducing the catabolism of liver glycogen with an ensuing rise in blood glucose. Our blood biochemistry test results of 9.5 month old Gbe^ys/ys^ mice demonstrate that upon treatment with 144DG11, the characteristic hypoglycemia and hyperlipidemia of the mice were corrected to control levels (Fig. 3G). Muscle (creatine kinase) and liver (alanine transferase) functions were not affected by this treatment (Fig. 3G).

### *144DG11* enhances catabolism in glycogen overloaded APBD patient cells

The RQ shift towards carbohydrate catabolism observed in vivo prompted us to investigate whether carbohydrate catabolism is also up-modulated intracellularly. To that end, and especially since glycogen levels are highly variable among fibroblasts derived from different APBD patients (Fig. 4A), we first aimed at inducing a physiological glycogen overload, or glycogen burden, condition, equivalent to the one found in tissues. We found that glycogen burden can be produced by 48 h glucose starvation followed by replenishment of the sugar for 24 h, which possibly induces accelerated glucose uptake with ensuing glycogen synthesis. This starvation/replenishment condition indeed increased intracellular glycogen levels, as demonstrated by PAS staining (Fig. 4B). Furthermore, a multiparametric high-content imaging-based phenotyping analysis revealed that under glycogen burden conditions, cell area, nuclear intensity and, importantly, mitochondrial mass features (see boxes in Fig. 4C) deviated from healthy control (HC) more than glucose starved-only cells did. Therefore, we selected this glycogen burden condition to analyze catabolism at a cell level using Agilent’s Seahorse ATP Rate Assay. Our results (Fig. 4D) show that at the cell level, 144DG11 increased not only overall ATP production, but also the relative contribution of glycolytic ATP production at the expense of mitochondrial (OxPhos) ATP production. This phenomenon was observed in both HC and APBD patient skin fibroblasts. Acute on assay supplementation of 144DG11 was more effective at augmenting the glycolytic contribution to ATP production than 24 h pretreatment with the compound. These results suggest that glucose derived from the 144DG11-mediated enhanced carbohydrate catabolism is exploitable for ATP production.

**Fig. 4.**
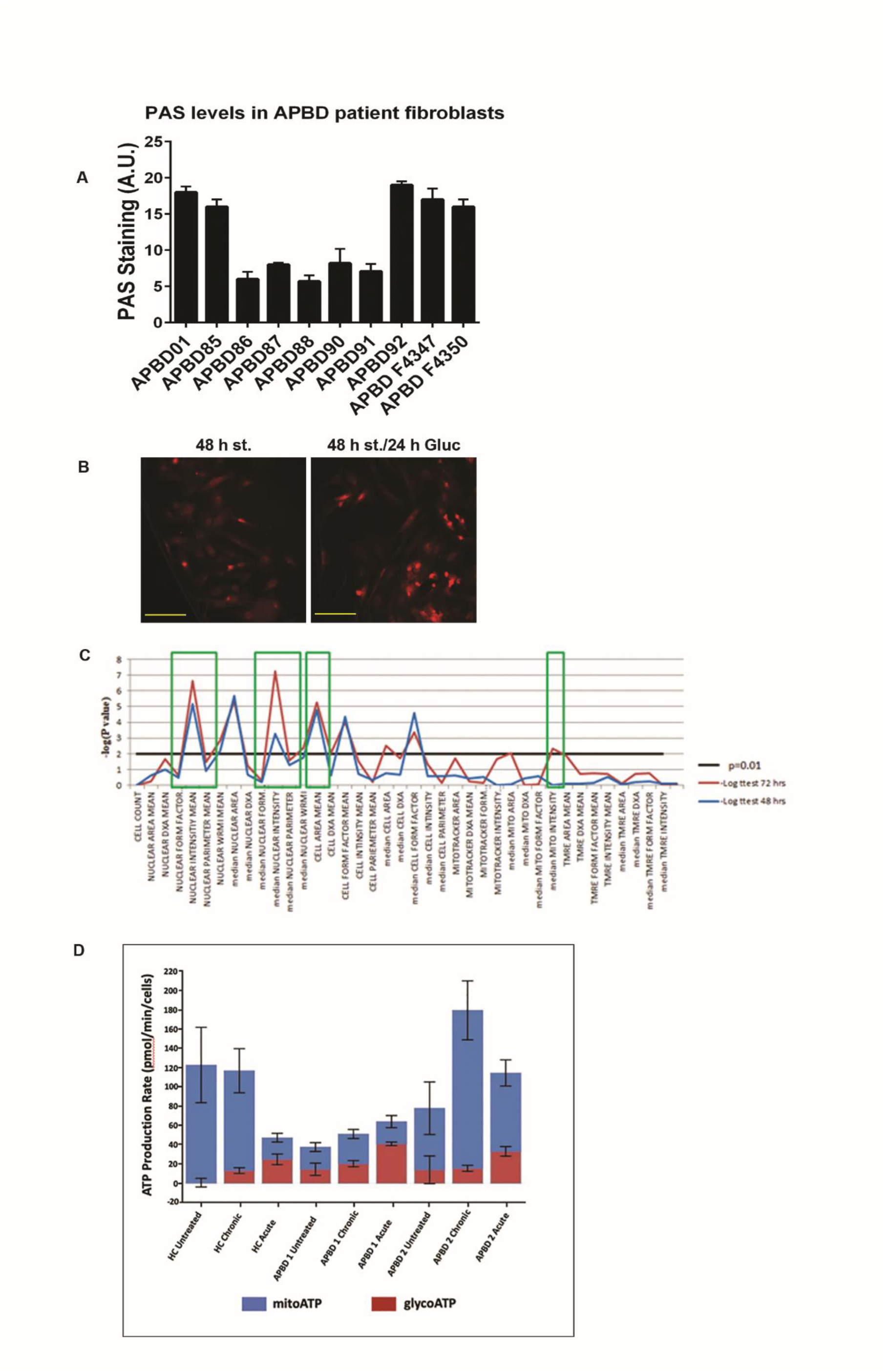
Glycogen burden in APBD fibroblasts and effect of 144DG11 on ATP production. **(A)** PAS staining for total glycogen in skin fibroblasts from different APBD patients. Imaging done by InCell2200 (see Methods). Means are statistically different from each other (p<0.0001, One Way ANOVA). **(B)** PAS staining for total glycogen in APBD87 fibroblasts glucose-starved for 48 h (*left*), or glucose starved and then replenished for the last 24 h to induce glycogen burden (*right*). Image acquisition was done by Nikon Eclipse Ti2 microscope using a 40x PlanFluor objective and CY3 filter. Scale bar, 100 µm. **(C)** Image-based multiparametric phenotyping (see Supplementary Materials) of APBD fibroblasts under 48 h glucose-starvation (blue), or starvation and glucose replenishment as in **(B)** (red). Black line, level of significance p=0.01. **(D)** Glycolytic (red) and mitochondrial (blue) ATP production determined by Agilent’s Seahorse machine and ATP rate assay kit. HC and APBD patient fibroblasts were serum/glucose-starved for 48 h and then full medium was replenished for 24 h without (untreated), or with (chronic) 50 µM 144DG11. Acute, 50 µM 144DG11 were added on assay for 20 min after 24 h of serum/glucose replenishment. Readings were normalized to cell number as determined by Crystal Violet staining. Shown are mean and SD values based on n=6 repeats. In all experiments (HC, p<0.002; APBD1, p<0.0005; APBD2, p<0.1, One Way ANOVA with Sidak’s post-hoc correction for multiple comparisons) glycolytic ATP production was increased, as compared to untreated control, by acute supplementation of 144DG11. Chronic supplementation of 144DG11 increased glycolytic ATP production only in HC (p<0.017). Mitochondrial ATP production was reduced by acutely supplemented 144DG11 in HC (p<0.002) and increased by chronically supplemented 144DG11 in APBD1 (p<0.05) and APBD2 (p<0.01).

### *144DG11* binds to the lysosomal membrane protein LAMP1

We next investigated the mechanism of action of 144DG11. To that end, we first decided to determine its molecular target. Nematic protein organization technique (NPOT, Inoviem, Ltd.) was applied to homogenates of APBD patient fibroblasts. The NPOT analysis revealed protein hetero-assemblies uniquely generated around 144DG11 only when it was added to the cell homogenates (Fig. 5A). The next step in this analysis identified the interactome of protein targets interacting with 144DG11 in APBD patients’ fibroblasts. Interestingly, as revealed by Inoviem’s gene ontology analysis based on several bioinformatic tools, proteins in the hetero-assembly interacting with 144DG11 in APBD patient fibroblasts are autophagy, or lysosomal proteins (Fig. 5B). Moreover, we tested by cellular thermal shift assay (*16*) the specific interaction of 144DG11 with 6 of the 8 targets discovered by NPOT. Our results (Fig. 5C) suggest that LAMP1, and not other protein targets, directly interacts with 144DG11. This finding is related to a novel pathogenic hypothesis connecting cellular glycogen overload with glycogen trafficking to lysosomes via Starch Binding Domain containing Protein 1 (*17*). To validate 144DG11’s interaction with LAMP1, we used surface plasmon resonance (SPR) technology. Our SPR data (Fig. 5D) show a specific and dose-dependent binding of 144DG11 to the luminal portion of LAMP1 only at the lysosomal pH 4.5-5 and not at the cytoplasmic pH 7, with some binding starting at the intermediate pH 6. Taken together, these results constitute a strong and acceptable evidence that the specific target of 144DG11 is the type 1 lysosomal protein LAMP1, widely used as a lysosomal marker and a known regulator of lysosomal function. However, the apparent K_D_ of this binding was relatively high (6.3 mM), which we ascribed to the slow k_on_ (rate of association in Fig. 5D, pH 4.5). We hypothesized that this slow rate of association could be explained by inhibited diffusion of 144DG11 due to the bulky oligosaccharides at the glycosylation sites. Therefore, we repeated the SPR experiments with a chemically deglycosylated luminal LAMP1 domain. Indeed, deglycosylated LAMP1 bound 144DG11 with an apparent K_D_ of 52.5 nM., possibly due to profound structural changes induced by the deglycosylation (Fig. 5E). We further investigated 144DG11 binding to LAMP1 by structure-based computational docking. In the search for a putative binding site for 144DG11 in LAMP1, we analyzed the N- and C-terminal subdomains of its luminal domain (residues A29R195 and S217-D378, respectively), which have a similar topology (*18*) . These domains were modeled (Supplementary Materials) at the intralysosomal pH 5 based on the known crystal structure of mouse LAMP1 C-terminal domain (PDB ID 5gv0). Possible binding sites were identified by three different computational tools: SiteMap, FtSite and fPocket, and docking computations of 144DG11 and a set of decoy molecules were performed to every putative binding site by the Glide algorithm. Fig. 5F shows the 144DG11 LAMP1 putative binding pocket (residues F50-D55, N62, L67, F118, Y120-L122, T125, L127-S133, N164-V166), predicted by all three tools and with high selectivity to 144DG11 according to the docking results. Prediction of the same binding site by three different programs is very rare and thus strongly suggests that 144DG11 binds to the specified site at the N-terminal of LAMP-1. As can be seen in Fig. 5F, Asn-linked oligosaccharides face away from the predicted 144DG11 binding site and are therefore not expected to directly interfere with its binding. However, they might still affect 144DG11 diffusion in agreement with the significantly reduced K_D_ in deglycosylated LAMP1 (Fig. 5E).

**Fig. 5.**
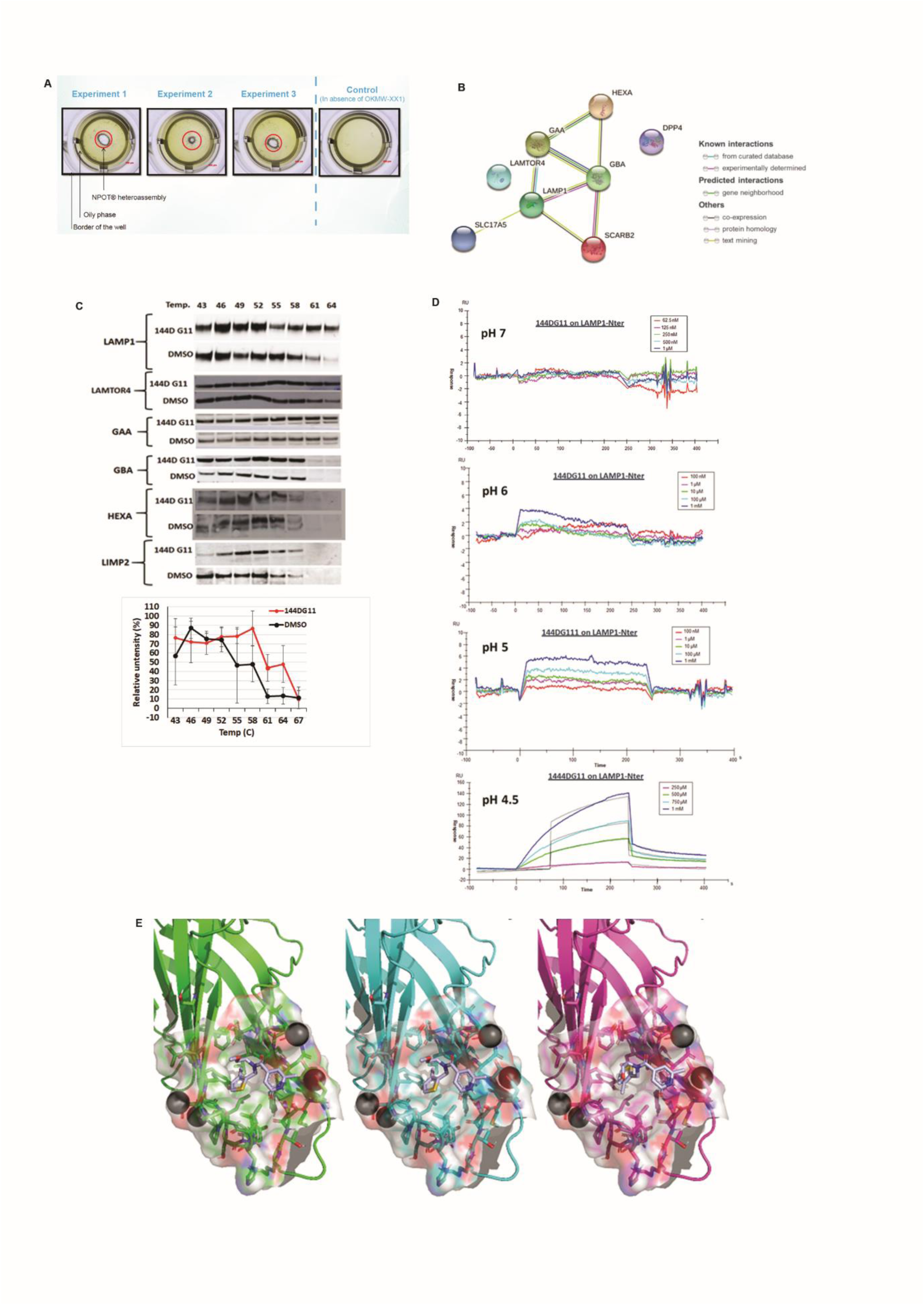
144DG11 interacts with LAMP1 and its associated interactome. **(A)** Hetero-assembly forms around 144DG11 and not around endogenous molecules as shown by the liquid crystals formed in experiments 1-3 (see Supplementary Materials). **(B)** STRING network of targets at the interactome of 144DG11. Targets also show strong protein-protein interactions among themselves, as connectors show (see legend). SLC17A5, Solute Carrier Family 17 Member 5; LAMP-1, Lysosomal Associated Membrane Protein 1; LAMTOR4, Late endosomal/lysosomal adaptor, MAPK and MTOR activator 4; GBA, Glucosylceramidase; SCARB2 (aka LIMP2), Scavenger Receptor Class B Member 2; DPP4, Dipeptidyl peptidase-4; GAA, Lysosomal alpha-glucosidase; HEXA, Beta-Hexosaminidase Subunit Alpha. **(C)** Cellular thermal shift assay (CETSA) of different targets of the 144DG11hetero-assembly **(B)**. Cell lysates were treated with 50 uM compound A, or an equivalent volume of DMSO and then heated to the indicated temperature to generate a melt curve. Precipitated protein and cellular debris were separated by centrifugation and the soluble fraction was subjected to SDS-PAGE and immunoblotted with the indicated Abs (see (*16*) for details). Only LAMP1 was significantly protected by 144DG11 from heat-mediated denaturation (*i.e.*, its Tm was increased). Consequently, it was concluded that only LAMP1 specifically interacted with 144DG11. **(D**) Surface plasmon resonance assays to test 144DG11 binding to LAMP1. Luminal N terminus (aa 29-382) of bioactive LAMP1 (binds Galectin-3 with apparent K_D_ < 25nM) with post-translational modifications (R&D systems (4800-LM-050)) was reconstituted at 200 µg/mL in PBS and immobilized to a gold-carboxymethylated dextran sensor chip by direct amine coupling. Sensogram experiments consisting of association and dissociation at the indicated concentration ranges and pH values were then conducted. Results show that dose-responsive association of LAMP1 to 144DG11 started at pH 6, was partial at pH 5 and was clearly demonstrated at the lysosomal pH 4.5-5. All sensorgrams correspond to normalized signal (*i.e*., reference surface and buffer signals were substracted). **(E)** *Upper panel*, LAMP1 was deglycoslated as detailed in Supplementary Materials. RNase B is a positive control for a glycosylated protein. Glycosylation status is shown after short (24 h) or long (72h) dialysis. The results demonstrate a full deglycosylation of both LAMP1 luminal part and RNase B, with unique bands appearing after a long dialysis. *Lower panel*, Surface plasmon resonance, performed as in **(D)** of deglycosylated LAMP1 luminal domain. **(F)** Three binding modes of 144DG11 (gray) according to LAMP1 grids that were predicted by SiteMap (green), fPocket (cyan) and FtSite (magenta). Two out of three binding modes (SiteMap and fPocket) are identical. In FtSite part of the molecule went through a rotation relative to the other two. See Supplementary Materials.

### *144DG11* enhances LAMP1 knockdown-induced autolysosomal degradation and catabolism of glycogen

144DG11 increased autophagic flux in APBD primary fibroblasts. This is demonstrated by an increased sensitivity to lysosomal inhibitors in the presence of 144DG11. As can be seen in Fig. 6A, lysosomal inhibitors increase the LC3ii/LC3i ratio (autophagic halt) more in 144DG11 treated than in untreated cells. Increase in autophagic flux by 144DG11 was also illustrated by lowering the level of the autophagy substrate p62 (Fig. 6A). Moreover, transmission electron microscope analysis of liver sections of the APBD modeling Gbe^ys/ys^ mice demonstrates a decrease in lysosomal glycogen following treatment with 144DG11 (Fig. 6B).

**Fig. 6.**
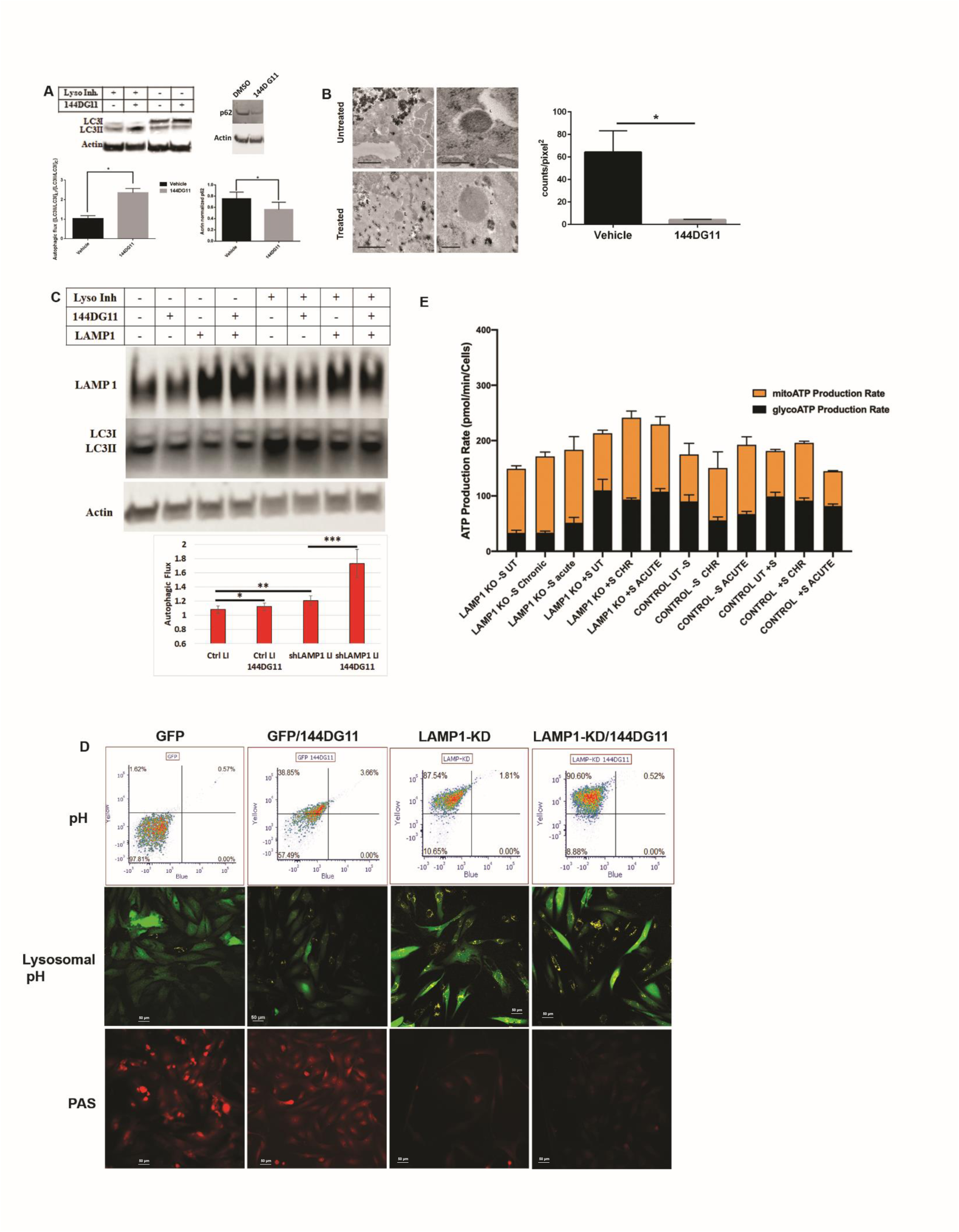
LAMP1-KD and 144DG11 enhance autophagic flux. **(A)** Autophagic flux, determined by the extent of lysosomal inhibitors-dependent increase in the ratio of lipidated to non-lipidated LC3 (LC3II/LC3I), is significantly increased by 144DG11 (50 µM, 24 h). *Upper left panel,* a representative blot. *Lower left panel,* densitometric quantification of autophagic flux showing significant increase by 144DG11 (n=4, p<0.0023, Student’s t-test). *Upper right panel*, 144DG11-mediated increase in autophagic flux is also demonstrated by enhanced degradation of the autophagy substrate p62. A representative blot is shown. *Lower right panel,* densitometric quantification of p62 levels showing significant decrease by 144DG11 (n=5, p<0.0099, Student’s t-test). **(B)** Representative TEM images of liver tissue from 9.5 month old Gbe^ys/ys^ mice treated with 144DG11, or 5% DMSO vehicle. G, Glycogen (alpha particles) and polyglucosan (structures with variable electron densities), L, Lysosomes, M, Mitochondria. Note accumulation of glycogen and polyglucosan in lysosomes corrected by the treatment (right). The low magnification samples (left) show the overall higher levels of glycogen and polyglucosan in samples collected from untreated *v* 144DG11 treated animals. *Right panel*, lysosomal glycogen particles were quantified by ImageJ “count particle” tool. Treatment led to a significant reduction in lysosomal glycogen (n=4, *p<0.03, Student’s t-test). **(C)** LAMP1 knockdown increases autophagic flux, which is further facilitated by 144DG11. *Upper panel*, LAMP1 knocked down and control APBD primary skin fibroblasts were treated or not with 144DG11 and lysosomal inhibitors (LI). Imunoblotting of cell lysates with LAMP1, LC3 and actin abs shows that autophagic flux (calculated as in **(A)**) is increased by LAMP1 knockdown and further by 144DG11. *Lower panel*, densitometric quantification of autophagic flux (n=3, *, p<0.1; **, p<0.05; ***, p<0.01 (Student’s t-tests)). **(D)** Lysosomal pH changes determined in APBD primary fibroblasts transduced with lentiviruses encoding for GFP or GFP-shLAMP1 and treated or not with 144DG11 for 24 h. *Upper panel,* Transduced cells were stained with 5µM LysoSensor Yellow/Blue DND-160 just before acquisition in a BD-LSRII flow cytometer using the 375 nm laser line and the 505 LP and 450/50 BP filters to detect the yellow and blue emissions of Lysosensor, respectively. The 488 nm laser line with the 530/30 BP filter was used to detect GFP fluorescence. Yellow Lysosensor and GFP fluorescence were compensated. N.B., yellow to blue median fluorescence ratio (Y/B) positively correlates with acidification. 144DG11 very slightly acidified GFP cells (Y/B(GFP/144DG11)>Y/B(GFP), p<0.12), but significantly acidified LAMP1-KD cells (Y/B(LAMP1-KD/144DG11)>(Y/B(LAMP1-KD), p<0.03). The most significant acidification was caused by LAMP1-KD (Y/B (LAMP1-KD)>Y/B(GFP), p<0.007). n=3, Student’s t-tests. *Middle panel*, cells treated with Lysosensor were analyzed by the Nikon A1R confocal microscope using the 488 nm and 405 nm laser lines for exciting GFP and Lysosensor (only yellow fluorescence), respectively. *Lower panel*, the same cells as indicated were stained for PAS as in Fig. 4B. **(E)** ATP production rate assay in LAMP1-KD and GFP (Control) cells treated for 24 h (chronic) or on assay (acute) with 144DG11. See Fig. 4B for details and text for statistical analysis (multiple t-tests, One Way ANOVA with Sidak’s post-hoc corrections).

To determine the functional importance of the interaction between 144DG11 and LAMP1, we knocked down the latter using a lentiviral vector carrying GFP tagged shRNA against LAMP1. As LAMP1 knockdown (KD) becomes cytotoxic 24 h post expression (or 96 h post lentiviral infection), LAMP1-KD experiments in Figs. 6C and 6D were conducted under 24 h serum starvation condition, without glucose replenishment (Fig. 4), to both induce autophagy and maintain cell viability. We expected LAMP1-KD to neutralize the effect of 144DG11 allegedly mediated by its interaction with LAMP1. Surprisingly, however, supplementation of 144DG11 to LAMP1 knocked down (*N.B.*, not knocked out) cells enhanced the knockdown effect: Autophagic flux, enhanced by LAMP1-KD, was further enhanced by the LAMP1 interacting 144DG11 (Fig. 6C). The observation that 144DG11 enhances LAMP1-KD effect suggests that the interaction of 144DG11 with LAMP1 is inhibitory, as many other small molecule-protein interactions are. Furthermore, to test whether LAMP1-KD and 144DG11 enhanced autophagic flux by improving lysosomal function, we quantified lysosomal acidification using the pH ratiometric dye Lysosensor, which quantifies pH based on the yellow/blue emission ratio. Our results show that both LAMP1-KD and 144DG11 treatment (in GFP and LAMP1-KD APBD cells) led to lysosomal acidification, but more so LAMP1-KD. As our confocal microscope is missing the 360 nm excitation line, we were unable to detect blue fluorescence microscopically. Therefore, we show by flow cytometry (Fig. 6D, upper panel) the overall cellular acidification as an increase in 375 nm-excited yellow/blue emission, and by confocal microscopy that this acidification is associated with brighter yellow fluorescence in lysosomes (Fig. 6D, middle panel). Importantly, as demonstrated by PAS staining, LAMP1 knockdown also reduced cellular glycogen levels, an effect which was slightly enhanced by 144DG11 in APBD fibroblasts transduced with both GFP control and shLAMP1-GFP lentiviruses (Fig. 6D, lower panel).

To test the effect of LAMP1-KD and 144DG11 on fuel utilization, we again used the ATP Rate Assay (Fig. 4) in LAMP1-KD and control APBD fibroblasts acutely or chronically treated with 144DG11. Our results (Fig. 6E) show that starvation was more restrictive (lowered overall ATP production) in LAMP1-KD (LAMP1-KD-S UT *v* LAMP1-KD+S UT, p<0.0001) than in GFP-transduced controls (Control UT-S *v* Control UT+S, p<0.36). In LAMP1-KD cells, starvation also increased the relative contribution of respiration to ATP production (78% in LAMP1-KD-S UT *v* 48% in LAMP1-KD+S UT (orange bars)). These observations are in line with the higher ATP production efficiency of respiration as compared to glycolysis and with a possibly higher ATP demand of LAMP1-KD, as compared to control cells, as suggested by their higher overall ATP production rate in basal conditions (*cf.* LAMP1-KD+S UT with Control+S UT, p<0.01). The effect of 144DG11 on LAMP1-KD and control cells was in accordance with its selective increase of catabolic (ATP generating) autophagic flux in LAMP1-KD cells, as compared to control cells (Fig. 6C): In non-starved conditions, supplementation of 144DG11 significantly increased total and respiratory ATP production in LAMP1-KD cells (*cf*. LAMP1-KD+S UT with LAMP1-KD+S Chronic (p<0.03 for total, p<0.0008 for respiratory) and LAMP1-D+S Acute (p<0.01 for total and respiratory)), while it only slightly influenced ATP production, and even acutely decreased it, in control cells (*cf*. Control UT+S with Control+S Chronic (p<0.1) and Control UT+S Acute (p<0.0008 for decrease)). Under starved conditions, control cells only increased respiratory ATP production in response to the transient effects of acutely supplemented 144DG11 (*cf*. Control UT-S with Control-S Acute, p<0.004). No significant effect of chronic supplementation of 144DG11 was observed in control cells (*cf*. Control UT-S to Control-S Chronic, p<0.3). In contrast, starved LAMP1-KD cells increased both respiratory and glycolytic ATP as a response to acute supplementation of 144DG11, possibly reflecting short-term diversion of glucose derived from glycogen degradation to glycolysis (*cf*. LAMP1-KD-S UT with LAMP1-KD-S Acute (p<0.0003 for glycoATP, p<0.003 for mitoATP). In response to chronically administered 144DG11, only respiratory ATP production increased (*cf.* LAMP1-KD-S UT with LAMP1-KD-S Chronic (p<0.15 for glycoATP, p<0.0002 for mitoATP) in LAMP-KD cells.

### *144DG11* reduces lysosomes and restores aberrant mitochondrial features

As we showed that the mode of action of 144DG11 involves lysosomal catabolism which increases ATP production, we decided to investigate whether the cellular features modulated by 144DG11 are relevant to its catabolic effects. As a first step, we determined that APBD and HC fibroblasts are phenotypically different in general and that the APBD phenotype is significantly more heterogeneous (Fig. 7A). This was done using image-based high content analysis (HCA) of thousands of cells per sample in the InCell 2200 image analyzer, followed by principal component analysis. 45 cellular features from 17 APBD and 5 age matching HC fibroblasts were analyzed under environmental control conditions. Our findings suggest plasticity of the affected phenotype and thus predisposition to interventional restoration (*e.g.* by 144DG11). As 144DG11 targeted LAMP1 and lysosomal and catabolic functions (Figs. 4-6), we focused on its effects on lysosomes and mitochondria, as analyzed by imaging and bioenergetic characterization. Fig. 7B shows that 144DG11 reduces lysosomal staining intensity and area under starvation. This lysosomal reduction, observed also in healthy as compared to lysosomal impaired cells (*19*), might be associated with the compound’s mediated improvement of autophagic flux, induced in starvation, and lysosomal function (Fig. 6). Mechanistically, lysosomal reduction by 144DG11 could be mediated by its effect on LAMP1, a major component of the lysosomal membrane which maintains its integrity. Fig. 7C shows that 144DG11 has also hyperpolarized the mitochondrial membrane potential (MMP), depolarized by the diseased state in APBD, and increased mitochondrial biomass, also reduced by the diseased state. Both results are in accordance with possibly increased mitochondrial fueling by the enhanced autophagic catabolism. To test this possibility, we conducted a bioenergetic analysis of the effect of 144DG11 in glycogen burden conditions, demonstrated to enhance its effects (Fig. 4). Our results (Figs. 7D and S8) show that 144DG11 has increased basal and maximal respiration, as well as mitochondrial ATP production, coupling efficiency and capacity to offset proton leak, or respiratory control ratio (state 3/state 4 respiration).

**Fig. 7.**
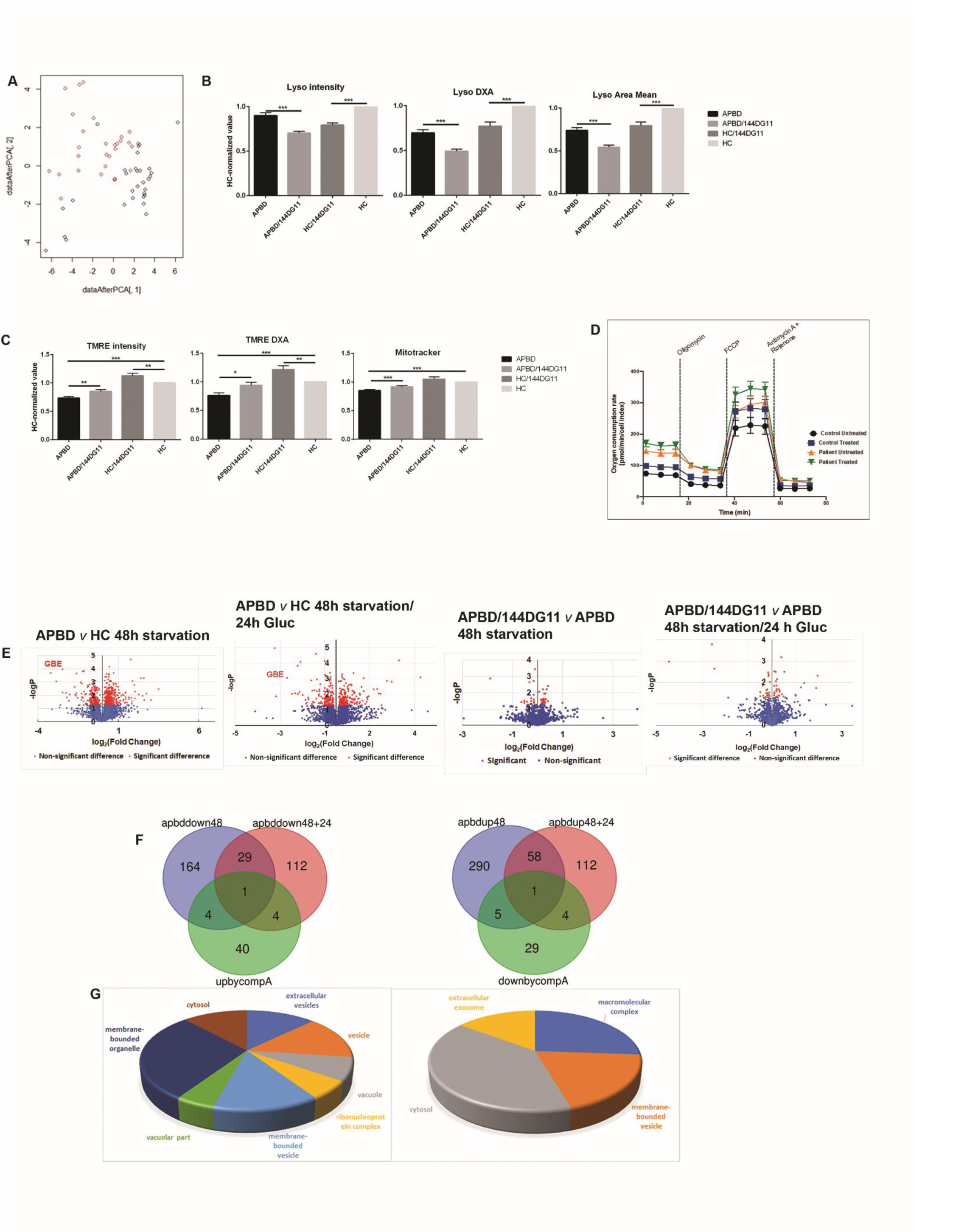
Phenotypic and proteomic effects of 144DG11. **(A)** Principal Component Analysis of IBP parameters in HC (black circles) and APBD (red circles) fibroblasts. **(B)** Morphological characterization of lysosomes based on the fluorescent lysotracker staining of n=4 HC and n=4 APBD patient skin fibroblasts starved for 48 h and treated or not with 50 uM 144DG11 for 24 h. Values are presented as deviations from untreated HC. 144DG11 has reduced lysotracker mean intensity, integrated intensity (DXA), and area in both HC and APBD fibroblasts (p<0.0001, One-Way ANOVA with Sidak’s multi-comparison post-hoc test). **(C)** Functional (mitochondrial membrane potential reported by TMRE mean and integrated intensity) and biomass (mitotracker) characterization of mitochondria in HC and APBD cells treated and analyzed as in **(B)**. Mean (p<0.0001) and integrated (p<0.0006) TMRE intensities, as well as mitochondrial biomass (p<0.0001), were reduced in APBD *v* HC. 144DG11 increased TMRE mean (p<0.009, p<0.006) and integrated (p<0.01, p<0.001) intensities in APBD and HC cells, respectively. Mitotracker intensity was increased by 144DG11 only in APBD cells (p<0.0001). **(D)** Oxygen consumption rate (OCR) in HC and APBD patient cells starved for 48 h and then treated with 50 uM 144DG11 for 24 h in full medium. Cells were analyzed by Agilent’s Seahorse machine and Mitostress kit. Shown is a representative experiment (n=3 biological replicates) based on n=6 technical replicates. 144DG11 treatment significantly increased basal and maximal respiration and coupling efficiency in both HC and APBD fibroblasts and ATP production only in APBD fibroblasts (Fig. S8). **(E)** Volcano plots of the proteins affected by APBD and 144DG11 under starvation and glycogen burden conditions. Protein extracts were made for LC/MS proteomics analysis from APBD (n=3) and HC (n=3) cells cultured for 48 h under either glucose and serum starvation or for additional 24 h in complete medium with either 50 uM 144DG11 or 5% DMSO vehicle. Red, significantly modulated proteins. **(F)** Venn diagrams of proteins down-modulated by APBD and up modulated by 144DG11 and *vice versa* under starvation (48) and glycogen burden (48+24) conditions. **(G)** Gene ontology of proteins up-modulated (left) and down-modulated (right) by 144DG11.

To investigate the global therapeutic impact of 144DG11, we analyzed the effect of the disease and the treatment on protein expression under different conditions (Data file S1). As shown in Fig. 7D, under 48 h starvation 12.2% and 6.8% of the 2,898 proteins analyzed were respectively up and down modulated in APBD-patient as compared to HC cells. Interestingly, endocytosis, a pathway implicated in lysosomal biogenesis and function, is a KEGG pathway upmodulated in APBD cells (Fig. S9), while oxidative phosphorylation is down-modulated in APBD cells (Fig. S10). As an important control, GBE was indeed down-modulated in the APBD cells (Fig. 7D). When starvation was followed by glucose supplementation (glycogen burden, Fig. 4), only 6% of the proteins were up-modulated and 5% down-modulated, possibly suggesting a more specific subset of proteins was required for managing the excess glycogen burden. For instance, autophagy proteins (Fyco1, Rab12, Rab7A, PIP4K2B, SQSTM1, and SNAP29) were only up-modulated in APBD cells following glycogen burden (Data file S1). We then investigated the proteomic effect of 144DG11 in starved (48h starvation) and glycogen overladen (48h starvation/24h Gluc) APBD cells, which respectively modified only 1.7% and 1.3% of all proteins. The apparently corrective effect of 144DG11 can be uncovered by proteins down-modulated or up-modulated by the APBD diseased state, which were inversely up-modulated (Table 1) or down-modulated (Table 2) by 144DG11 (Fig. 7F). The discovered proteins (49 up-modulated, 39 down-modulated, Fig. 7F) were analyzed by the DAVID functional annotation tool according to the Cellular Component category, which included the highest number of proteins. Proteins up-modulated by 144DG11 belonged to 8 significant gene ontology (GO) terms, which included lysosomal, secretory pathways and oxidative phosphorylation proteins (Fig. 7G, left panel and Table 1) in accordance with the cell features modulated by the compound (Figs. 7B and C). Interestingly, proteins down-modulated by APBD and up-modulated (“corrected”) by 144DG11 (yellow, Table 1) were the lysosomal glycosylation enzymes iduronidase and phosphomannomutase2 under glycogen burden, whereas under starvation those were the nucleic acid binding proteins GRSF1 and HNRPCL1, apparently not directly associated with glycogen and lysosomal catabolism (yellow, Table 1). The lipogenetic protein HSD17B12 was decreased by APBD and induced by 144DG11 under both conditions (green, Table 1). Proteins down-modulated by 144DG11 belonged to 4 GO terms, which included secretory pathways and macromolecular complexes (Fig. 7G, right panel and Table 2). Proteins increased by APBD and contrarily reduced by 144DG11 belonged to lysosomal sorting (VPS16) and carbohydrate biosynthesis (NANS) in starved cells and to transcription (RUBL1), signal transduction (STAM2) and pH regulation (SLC9A1) in glycogen overladen cells. Interestingly, pharmacological inhibition of the Na^+^/H^+^ antiporter SLC9A1 induces autophagic flux in cardiomyocytes (*20*) as does its down-modulation in APBD fibroblasts by 144DG11 (Table 2). The protein down-modulated by 144DG11 in both starved and glycogen burden conditions is the retrograde traffic regulator VPS51 (Table 2, green) also implicated in lysosomal sorting. In summary, the APBD correcting effects of 144DG11 are at least partially related to lysosomal function whose modulation by the compound is well characterized here (Figs. 5, 6).

**Table 1:**
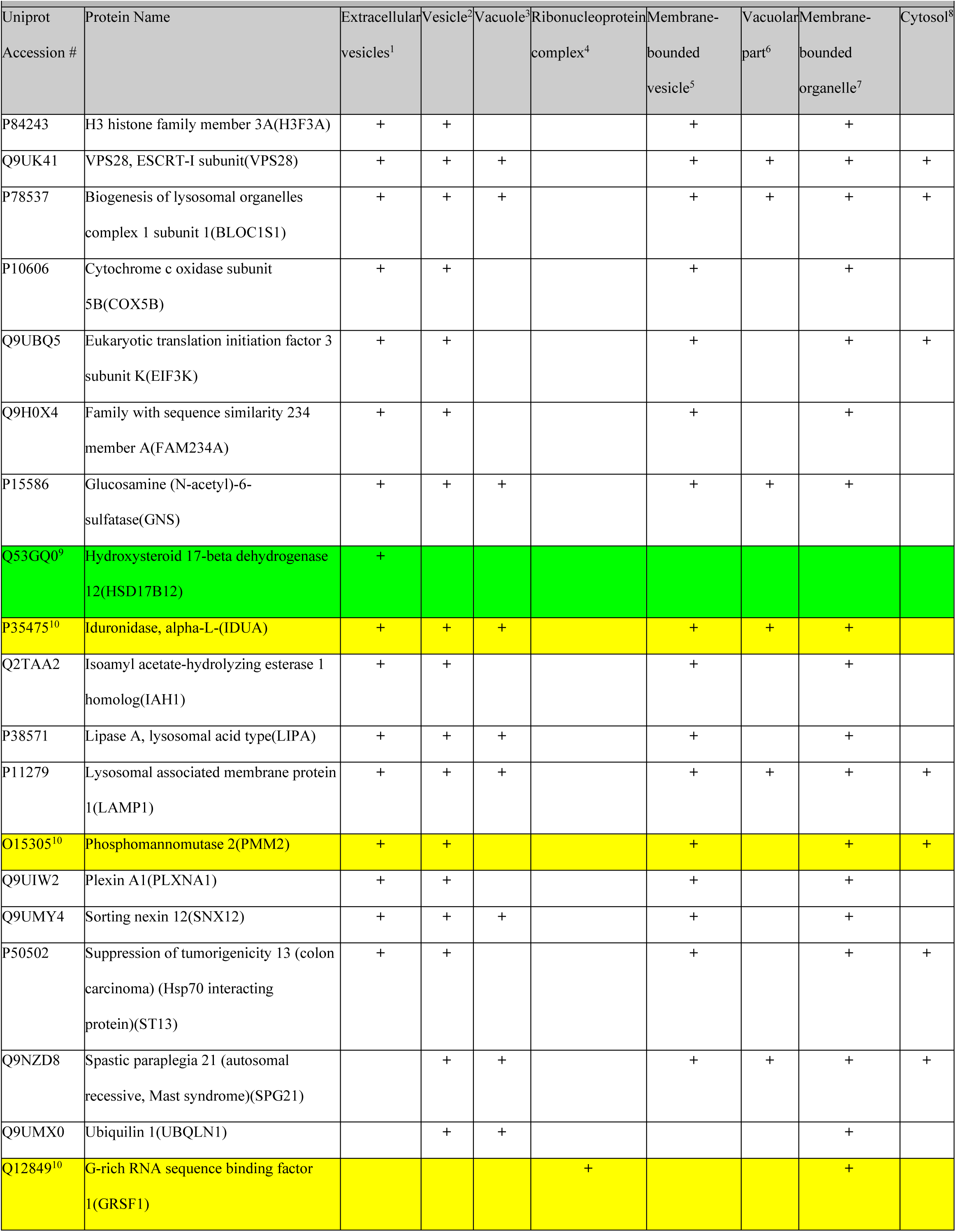
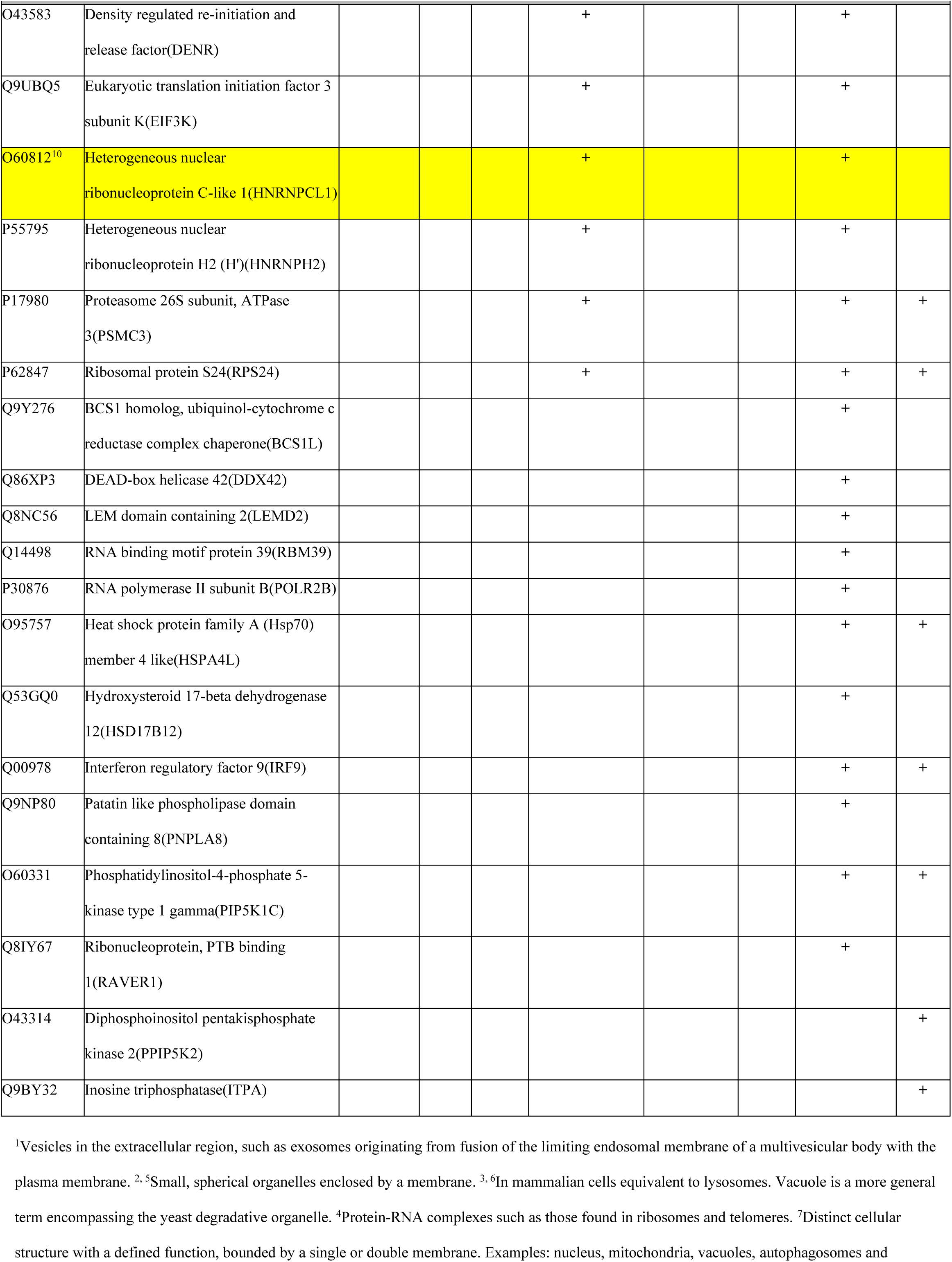
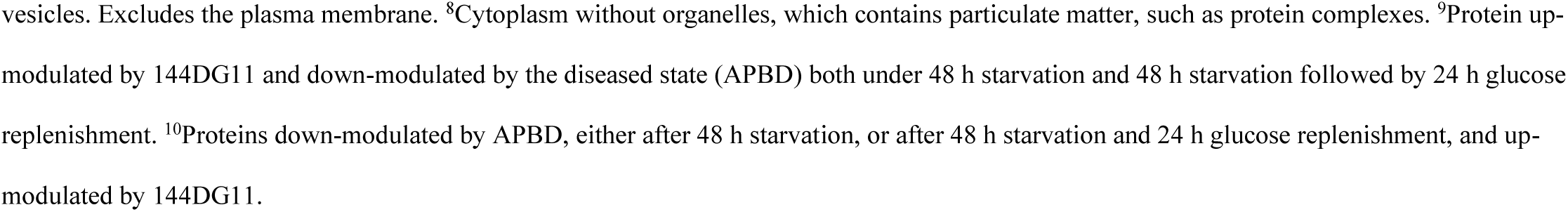
*Homo sapiens* proteins up-modulated by 144DG11

**Table 2:**
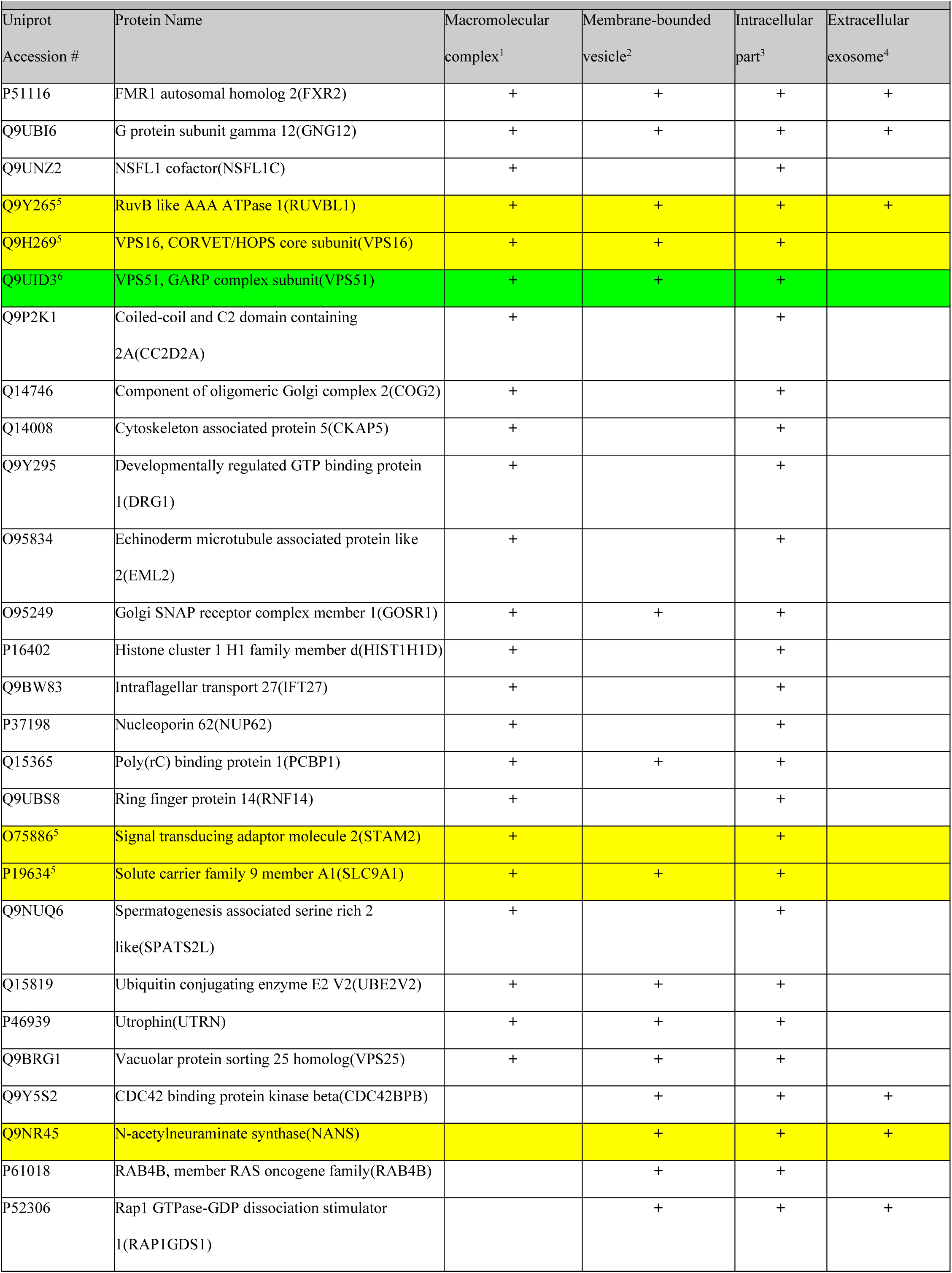
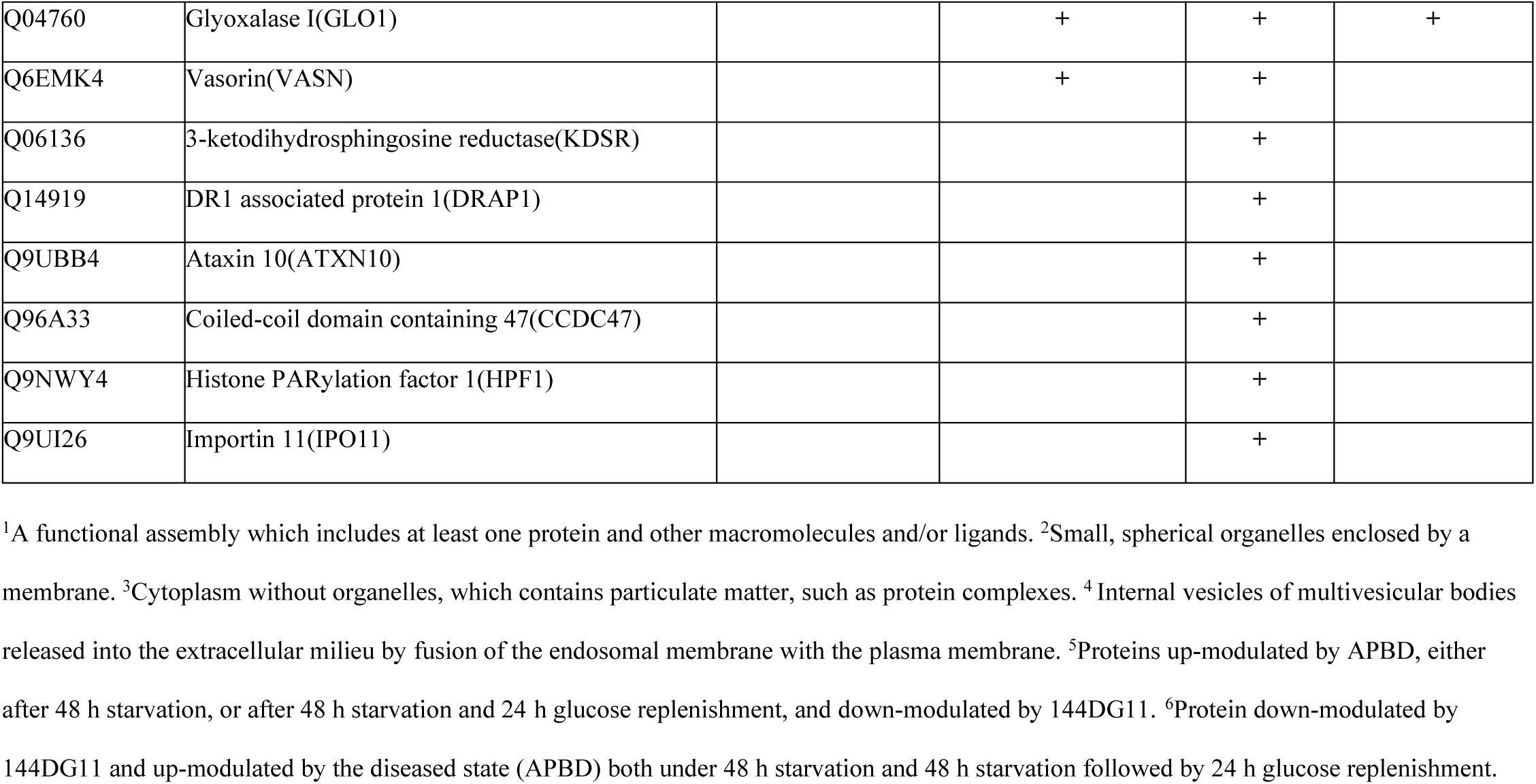
*Homo sapiens* proteins down-modulated by 144DG11

## Discussion

This work shows that the HTS-discovered hit 144DG11 can remedy APBD in in vivo and ex vivo models. Following 144DG11 treatment, we observed improvements in motor, survival and histological parameters (Figs. 1 & 2). As APBD is caused by an indigestible carbohydrate, these improvements suggested that 144DG11 affected carbohydrate utilization and thus encouraged us to conduct in vivo metabolic studies (Fig. 3). To our knowledge, this is the first in vivo metabolic study in a GSD animal model. Since APBD mice store glycogen as insoluble polyglucosan, we tested by indirect calorimetry whether 144DG11 can influence the capacity of these animals to use alternative fuels (fat) instead of mobilizing glycogen. However, the increased RQ induced by 144DG11 suggested that instead of using fat, treated animals actually increased carbohydrate burn, or that 144DG11 increased carbohydrate catabolism. This conclusion was supported by the 144DG11-induced increases in total energy expenditure, ambulatory activity, and meal size - all in line with catabolic stimulation. Since Gbe^ys/ys^ mice and APBD patients store glycogen as insoluble and pathogenic polyglucosan, its catabolism constitutes a therapeutic advantage. Glycogen catabolism is also a preferred therapeutic strategy for the following reason: In theory, therapeutic approaches to APBD should target either PG formation, or degradation of preformed PG or glycogen (Fig. S5, approaches 1 and 2, respectively). PG formation depends on the balance between GYS and GBE activity – the higher the GYS/GBE activity ratio, the more elongated and less branched soluble glycogen would form, which would preferentially form PG, as compared to shorter chains (*21*). Degradation of pre-existing PG and glycogen (PG precursor), on the other hand, as done by 144DG11, is a more direct target and is expected to be more efficacious than inhibition of de novo PG formation, as done by the GYS inhibitor guaiacol (*15*), which spares pre-made detrimental PG. Indeed, in a study in LD-modeling mice, it was sown that conditional GYS knockdown after disease onset is unable to clear pre-existing and detrimental Lafora PG bodies (*22*).

A key challenge in drug discovery is the determination of relevant targets and mechanism of action of drug candidates. To that end, we have applied here Inoviem’s NPOT protein target identification approach. This technique, recognized as a leading tool for identifying protein targets of small molecules (*23*), and which identified several therapeutically relevant targets (*24–26*), identifies compound-target interactions within the natural physiological environment of cells. This means that the entity identified is not the target *per se*, as in other technologies, but the primary target with its signaling pathway, or functional quaternary network. Determination of the cellular pathway modulated by the test compound, as done for 144DG11, is important for formulation of other drugs to the same pathway, which can significantly upgrade therapeutic efficacy in due course in the clinic. Moreover, NPOT can also confirm the specificity of target binding by filtering out promiscuous binders, and excluding binding to negative controls (in our case, negative compounds in the HTS) and to endogenous ligands (Fig. 5A). Nevertheless, while by these criteria 144DG11 binding to LAMP1, and through it to its functional quaternary network (Fig. 5B), was specific and manifested dose response and lysosomal pH dependence in the SPR validation (Fig. 5D), its apparent LAMP1 binding K_D_ was relatively high (6.3 mM), which seemingly could be an impediment towards its clinical application. This issue can be coped with as follows: 1. The pharmacologically relevant finding is that 144DG11 specifically interacted with a lysosomal-autophagosomal interactome (Fig. 5B) and that it wasn’t toxic (Figs S1-S4, Table S1). This finding rules out non-specific interaction with putative off-targets, which is the main concern in low affinity (high K_D_) ligands, especially since this low affinity is probably a by-product of oligosaccharide steric hindrance (Fig. 5E). 2. A conventional approach to improve the affinity of low-affinity pharmaceutical candidates is based on medicinal chemistry. In GSDs, such an approach was used for increasing the affinity of GYS inhibitors (*27*). However, as opposed to GYSs, whose reduction is relatively tolerable (*12, 28*), the LAMPs belong to the house keeping autolysosomal machinery (Fig. 5B), whose inhibition can compromise perinatal viability, as does, for instance, LAMP1-KD without a compensatory rise in LAMP2 (*29, 30*). Therefore, a high affinity LAMP1 inhibitor might be toxic, as was LAMP1-KD to APBD fibroblasts (Fig. 6), and the low affinity of the LAMP1 inhibitor we discovered, 144DG11, may actually constitute a clinical advantage by mitigating the repression of a household function. 3. The discovery of the LAMP1 containing hetero-assembly (Fig. 5B) as a functional network, rather than a single protein target, opens a therapeutic modality based on autophagy modulation, which actually expands the therapeutic target landscape. Autolysosomal network was discovered not only in Fig. 5B, but also by image high-content analysis, in conjunction with bioenergetic parameters, possibly modified by autophagy-associated changes in fuel availability (Figs. 7B and C). Other supports for the relevance of this pathway as a 144DG11 target come from our proteomics data (Fig. 7D-G) and from the actual boost of autophagic flux by 144DG11 in cells (Fig. 6).

Mechanistically, LAMP1 is a type I lysosomal membrane protein which, together with LAMP2, plays a pivotal role in lysosome integrity and function (*29, 30*). Consequently, LAMP1, but more so LAMP2 (*31*), are also important for lysosomal involvement in the autophagy process. Therefore, LAMP1 knockdown is often associated with decreased autophagy (*32, 33*). However, in agreement with our results, other works show that LAMP1-KD actually increased autophagic function (*31, 34*), which was also shown for another transmembrane lysosomal protein TMEM192 (*35*). These apparent disparities probably depend on cell type, assay conditions, and even the definition of autophagy, as autophagic flux is not always defined by susceptibility to lysosomal inhibitors (*36*). To predict the molecular mechanism of action of 144DG11 on LAMP1, we used structural-computational techniques. Our computational results predict that the 144DG11 binding site is located at the LAMP1:LAMP1 interaction interface (Fig. S6A) (located at the N-terminal domain), and suggest that the compound inhibits inter-LAMP1 interaction. According to experimental data (*18*), truncation of the N-terminal domain of LAMP1 impairs LAMP1/LAMP1 and LAMP1/LAMP2 assembly, while truncation of the more mobile LAMP2 N-terminal domain leads to the opposite effect (Fig. S6B). Therefore, we may assume that LAMP1 N-terminal domain promotes LAMP1/LAMP1 and LAMP1/LAMP2 interactions and that inhibition of LAMP1/LAMP1 or LAMP1/LAMP2 interactions at the N-terminal domain, by 144DG11, would lower LAMP1 effective lysosomal membrane density. Thus, 144DG11 treatment can be hypothesized to be equivalent to LAMP1-KD, which might explain its enhancement of the LAMP1-KD effect. The slight increase (1.2 fold) in LAMP1 levels induced by 144DG11 (Table 1, Data file S1) probably reflects binding-mediated stabilization (Fig. 5C) and presumably does not significantly counteract 144DG11-mediated reduction in membrane density, suggesting that it increases glycophagy by the documented (*29, 30*) increase in LAMP2 in lysosomal membranes upon LAMP1-KD. LAMP2 was observed to enhance autophagosome-lysosome fusion (and thus autophagic flux) by interaction with the autophagosomal peripheral protein GORASP2 (*37*). Alternatively, spacing of the lysosomal membrane by LAMP1-KD/144DG11 may enable glycogen import to the lysosome (and consequent degradation) by the STBD1 protein (*17*). Importantly, lysosomal glycogen degradation takes place in parallel with its cytoplasmic degradation (*38*), and, specifically, in a GSDIV mouse model, which also models APBD in mice, overexpression of the lysosomal glycogenase α-glucosidase corrected pathology (*39*).

In summary, this work demonstrates 144DG11 as a novel catabolic compound capable of degrading PG and over-accumulated glycogen by activating the autophagic pathway. This study lays the groundwork for clinical use of 144DG11 in treating APBD patients who currently have no therapeutic alternative. Moreover, it positions 144DG11 as a lead compound for treating other GSDs through safe reduction of glycogen surplus.

## Materials and Methods

### Study design

This work combines in vivo, ex vivo and in vitro studies on the therapeutic potential of the newly discovered compound 144DG11 for treating APBD. In the in vivo section, we tested 144DG11 for its capacity to correct disease phenotypes in Gbe^ys/ys^ female mice. Two arms, 5% DMSO vehicle and 144DG11, of initially n=7-9 animals each were used. These numbers were demonstrated retrospectively to provide sufficient power because, based on the means and SD obtained, a power of 80% is already attained at n=5 animals/arm (*40*). Additional open field, gait, and extension reflex tests (Fig. 1E-H) also included a C57BL/6 wt control arm of n=9 animals. Animals were excluded from the experiment if weight was reduced by >10% between sequential weightings or by >20% from initiation. Sample size was slightly reduced over time due to death. 150 µL of 250 mg/kg 144DG11 in 5% DMSO were injected twice a week. Vehicle control was 5% DMSO. Injection was intravenous (IV) for the first month, followed by subcutaneous (SC) injection due to lack of injection space and scarring in animal tails. We initiated the injections either at the age of 4 months, two months prior to disease onset, assuming a preferred prophylactic effect, or at the 6 months age of onset for comparison. Treatment was continued until the age of 10 months. The effect of 144DG11 on various motor parameters was tested approximately every two weeks. At the end of these experiments, some of the mice were sacrificed by cervical dislocation and tissues from n=3 wt, n=7 Gbe^ys/ys^ vehicle-treated, and n=9 144DG11-treated mice were collected, sectioned, fixed and stained for diastase-resistant PG by PAS (Fig. 2). Tissue glycogen was determined biochemically as described (*15*). In addition, 144DG11 pharmacokinetic profile was determined by LC-MS/MS in serum and tissues derived from n=3 mice/time point. Experimenters were blinded to treatment allocation.

Ex vivo studies were done in APBD patient-derived skin fibroblasts and in liver sections from Gbe^ys/ys^ mice, as liver had the highest PG levels. In vitro studies were conducted in cell lysates. In vivo work was approved by the Hebrew University IACUC.

### Statistical analysis

In Fig. 1 the significance of overall trends was tested by Two-way ANOVA with repeated measures. This test determines how a response is affected by two factors: 144DG11 *v* control, which is given repeatedly (hence repeated measures) and duration of administration. The Bonferroni test was used to compare between 144DG11 and vehicle in a way which corrects for the multiple comparisons and is therefore very robust (since the threshold for determining significance at each time point is reduced in a manner inversely proportional to the number of comparisons). Consequently, most differences at specific time points became insignificant due to the increase in the number of comparisons and sometimes we chose to also show the data of multiple t-tests which do not correct for multiple comparisons. In Figs. 4D and 6E we used One Way ANOVA with Sidak’s post-hoc correction for multiple comparisons. Other statistical tests used were Student t-tests.

*Please see Supplementary Materials for other Methods*.

## Supporting information

Supplementary Materials revised

## Acknowledgments

We would like to thank Inoviem Scientific and Zakharia Manevich, Edward Berenshtein, Hava Glickstein, William Breuer and Hadas Segev-Yekutiel from the Hebrew University for their valuable help in experimental procedures.

## Author contributions

HV, KM, JD, MM, US, SW-A, AD, YR, BD, HE, SB, KM, AP designed and performed experiments; HR, JT and BM designed experiments and critically reviewed the manuscript; AL contributed clinical samples; OK and MW designed, supervised and wrote the manuscript.

