## Supplementary Materials revised for "A new drug candidates for glycogen storage disorders enhances glycogen catabolism: Lessons from Adult Polyglucosan Body Disease models"

**Supplementary Materials and Methods**

*Western immunoblotting*

Soluble protein factions were mixed with sample buffer and a reducing reagent (Life Technology) and warmed for 10 min at 70°C for protein denaturation. Samples were loaded on 10% Bis-Tris gels or 4–12% gradient gels (Invitrogen) with PrecisionPlus® Pre-stained Protein Standard protein ladder (Bio-rad) in MES SDS running buffer (Life Technology). Samples were run for 25-45 min at 200V in the Mini Gel Tank 21 apparatus (Life Technology). Protein transfer onto nitrocellulose membrane was preform in iBlot® 2 Gel Transfer Device (Life Technology). Membranes were then incubated for one hour in 5% BSA (bovine serum albumin) in Tris buffered saline with Tween 20 (TBST, pH=8) blocking solution for 1h. Western blot (WB) primary antibodies (see below) were diluted in WB blocking solution and then added for overnight incubation at 4 °C. Blots were probed with HRP conjugated secondary antibodies (see below) diluted in WB blocking solution for 1h at room temperature, followed by three TBST washes**.** To detect protein bands, ECL femto-kit (CYANAGEN) was used for development of membranes and signal detection was performed by the Amersham Imager 600 (Danyel Biotech). Quantification of band intensities was performed by densitometry analysis using ImageJ (Research Services Branch). *List of primary antibodies (all applied at 1:1000 titer)*: LC3 from rabbit (Cell signaling), HEXA from rabbit (Abcam), LAMTOR4 from rabbit (Cell signaling), LIMP-2 from rabbit (Abcam), GAA from rabbit (Abcam), GBA from rabbit (Abcam), Actin- HRP from mouse (Abcam), LAMP1 from rabbit (Abcam), p62 from rabbit (Abcam). *Secondary antibody:* HRP-conjugated donkey anti-rabbit and anti-mouse (Abcam).

*Lentiviral infection*

Lentiviral particles hosting LAMP1 shRNA fused to EGFP, or EGFP alone control plasmids were mixed with 8 µg/ml polybrene and supplemented to 80% confluent fibroblasts in fresh full medium for 8 h. Following incubation with lentiviral particles, medium was replaced and cells were examined by an epifluorescent microscope for EGFP fluorescence, which was normally observed after 72 h, as an indication of the expression of the lentiviral construct. Assays were conducted within 24 h after EGFP fluorescence was observed.

*Histological PG and glycogen determination*

Brain, heart, muscle, nerve fascicles (peripheral nerves), and liver tissues from wt and 144DG11 and vehicle treated $Gbe^{ys/ys}$ animals were separated to characterize the histopathological effects of 144DG11.

Tissues were extracted, fixed, embedded in paraffin, and sectioned. After deparaffinization, sections were treated for 5 min with 0.5% diastase to digest non-polyglucosan glycogen, leaving behind polyglucosan. Sections were then washed, stained for polyglucosan with PAS and counterstained with hematoxylin, and analyzed by light microscopy, all as described in (*15*). For biochemical glycogen determination, 100 mg of each tissue was subjected to alkaline hydrolysis and boiling followed by ethanol precipitation of glycogen. Glycogen was then enzymatically digested to glucose by amyloglucosidase (Sigma). Following digestion, total glycogen was determined based on the glucose content using the Sigma GAGO20 kit.

*Imaging and Image-based phenotyping*

APBD skin fibroblasts were seeded at 1,000 cells/well and cultured in specialized microscopy-grade 96-well plates (Grenier Bio-One, Germany). Following the different treatments, a mix of Thermo Scientific cellular fluorescent dyes in PBS was added to each well for 30 min at 37◦C in a 5% $CO_2$ incubator. This mix (Figs. 4C, 7B) included Hoechst (1 µg/ml, nuclear (DNA) stain), MitoTracker Green (500 nM, potential-independent mitochondrial stain), TMRE (500 µM, potential-dependent mitochondrial stain), and Calcein-AM Deep Red (0.5 µg/ml, cytosol stain). In Fig. 7C, only lysosomes were stained with LysoTracker Deep Red (75 nM). Cells were then fixed with 4% paraformaldehyde (PFA), washed with PBS and plates were transferred to an InCell2200 (GE Healthcare, U.K.) machine for image acquisition at 40× magnification. The output produced was based on comparative fluorescence intensity. Object segmentation was carried out using Multi-target analysis in the GE analysis workstation to identify the nuclei and cell boundary. All the assay parameters (including the acquisition exposure times, objective, and the analysis parameters) were kept constant for all assay repetitions. For PAS staining of glycogen (Figs. 4, 6C), fixed cells were washed with PBS, permeabilized with 0.1% Triton X-100, washed again stained and then imaged.

*Pharmacokinetics*

For pharmacokinetic analysis, 100 µL serum as well as brain, kidney, hind limb quad muscle, heart, liver, and spleen tissues were collected, homogenized and extracted with acetonitrile following established guidelines (*41*). Calibration curves were made with 0,1,10,100, and 1000 ng/ml 144DG11 in 1 mg/ml solutions of 4-tert-butyl-2-(4H-1,2,4-triazol-4-yl)phenol (ChemBridge) as internal standard (IS). Tissue samples were then dissolved in 1 mg/ml IS solutions and spiked with 0-1000 ng/ml 144DG11 to generate standard curves from which tissue levels of 144DG11 were determined. Samples were analyzed by the LC-MS/MS Sciex Triple Quad TM 5500 mass spectrometer.

*Target identification by nematic protein organization technique (NPOT)*

NPOT® was applied on human healthy fibroblasts and fibroblasts from two APBD patients. All the analyses were done by Inoviem Scientific Ltd. in a blinded manner. Protein homogenates from dry pellets of these fibroblasts were prepared by three cycles of fast freezing (liquid nitrogen) and slow thawing (on ice) and mixed at a maximal vortex speed for 30 seconds. Sample protein concentration was 50-66 mg/ml as determined by the BCA method. NPOT® is a proprietary technology offered by Inoviem Scientific dedicated to the isolation and identification of specific macromolecular scaffolds implemented in basic conditions or in pathological situations directly from human tissues. The technology is based on Kirkwood-Buff molecular crowding (*42, 43*) and aggregation theory (*44, 45*). It enables the formation and label-free identification of macromolecular complexes involved in physiological or pathological processes. The particular strength of Inoviem Scientific is the ability to analyze drug-protein and protein-protein interactions directly in human tissue, from complex mixtures without disrupting the native molecular conformation, consequently remaining in initial physiological or pathological condition.

Under laminar flow and sterile conditions, $10^{-6}$ M of compounds 144DG11 and a negative control from our HTS screen (*13*) were mixed separately with the protein homogenates (containing soluble and membrane proteins) and subjected to NPOT® isolation. The macromolecular assemblies associated with the ligand are separated using a differential microdialysis system, wherein the macromolecules (protein groups) migrate in the liquid phase based on their physico-chemical properties. The migrating macromolecules gradually grow from nematic crystals to macromolecular heteroassemblies thanks to the molecular interactions between the tested drug and its targets. The heteroassemblies were left overnight and isolated in a 96-well plate prior to identification by LC-MS/MS.

The formed heteroassemblies in presence of 144DG11 and the negative control in APBD-patients and HC fibroblasts are shown in Fig. S8. Each compound in contact with the indicated protein homogenates gave rise to clearly-defined heteroassemblies with common reticular morphology. The experiments were done in triplicate for each compound. For each of these biological replicates, heteroassemblies were isolated and their protein content analyzed by LC-MS/MS. The negative control is obtained with the protein homogenate in the NPOT® conditions without the addition of compound and does not present any aggregation. This further confirms that the formation of the heteroassembly is initiated by the compounds, and not by an endogenous small molecule, through their interactions with primary targets.

Under a Zeiss macroscope SteREO Discovery V8, each formed heteroassembly was isolated by microdissection and washed in acetone prior to solubilization in standard HBSS solution. Solubilized proteins were filtered through a 4-15% mini-PROTEAN gel. After migration, the gel was colored with a colloidal blue solution in order to visually estimate the number of proteins present in the gel, and the relative quantity of proteins to use for the following digestion step and injection in the LC-MS/MS instrument for proteomics analysis.

Proteomics was outsourced to the “Laboratoire de Spectrométrie de Masse Bio-Organique” (LSMBO) from the UMR 7178. Heteroassemblies were solubilized directly in 10 µL of 2D buffer (7 M Urea, 2 M Thiourea, 4% CHAPS, 20 mM DTT, 1 mM PMSF). Proteins were precipitated in acetate buffer and centrifuged for 20 minutes at 7500 g. Thereafter pellets were digested for 1 hour with Trypsin Gold (Promega) at 37°C. Trypsin Gold was resuspended at 1 µg/µL in 50 mM acetic acid, then diluted in 40 mM $NH_4HCO_3$ to 20 µg/mL. The samples were dried in Speed Vac® at room temperature. Peptides were purified and concentrated by using ZipTip® pipette tips (Millipore Corporation) before proceeding for mass spectrometry analysis through 1-hour nano-LC-MS/MS analyses protocol in an ESI-QUAD-TOF machine. Proteins were identified using Mascot software (Rank=1, score=25, minimal length=6 amino-acids, FDR=1%). For peptide mapping, the following database was used: - HumaniRTUN_DCpUN_JUS Bank (for human samples).

For data analysis and target deconvolution Inoviem Scientific developed its own database and software to allow an accurate and robust analysis of the proteins present in NPOT® datasets and simplify proteins ranking while removing protein contaminants. Inoviem Protein Ranking and Analysis (InoPERA®) database comprises all the NPOT® datasets obtained on various tissues, organs or cell lines, varied species and unrelated chemical compounds. InoPERA® software is then able to calculate the occurrence of one given gene in the entire database, or specific datasets matching defined criteria of species, organs etc. Inoviem removed contaminants that have been observed in NPOT® performed in human tissues and cells, which correspond to 613 NPOT® coupled LC-MS/MS analyses. Consequently, this tool is able to quickly highlight rare proteins within a dataset that would make new therapeutic targets (Fig. 5B).

Another bioinformatics resource - DAVID  was also used to find tissue-specific expression, gene-ontology and functional-related gene groups enrichment. Network enrichment within a dataset was investigated using STRING analysis (string-db.org). STRING is one of the core data resource of ELIXIR (as Ensembl or UniProt are) which contains known and predicted protein-protein interactions. Inoviem has used the stringent parameters, keeping only the known interactions ("experimentally determined" and "curated databases" interaction sources). This allowed deciphering the protein-protein associations within a complex dataset, which further completed the DAVID pathway analysis. In addition, Reactome (reactome.org) - a free, open-source, curated and peer-reviewed pathway database was used. This database provides intuitive bioinformatics tools for the visualization, interpretation and analysis of pathway knowledge to support findings obtained elsewhere.

In the bioinformatic pipeline, the first step of filtering consisted of removing the mass spectrometry "false positives", *i.e.,* the proteins found in one replicate and with only one specific peptide. Then, the datasets were compared in a 2 by 2 matrix (144DG11 and its respective negative control) in human skin fibroblast tissue. The next step of protein list analysis was identification of non-specific proteins, *i.e.* proteins that are found in a recurrent manner in all NPOT® experiments (InoPERA®). Contaminants (or "frequent hits") observed in human skin fibroblasts were removed. Cleared proteins lists of the interactome thus represent potential specific targets for 144DG11. Using this pipeline 28 proteins were found to interact specifically with 144DG11. 144DG11 interactome's specific protein lists were then analyzed independently by DAVID to find tissue-specific expression, gene-ontology and functional-related gene groups enrichment. The main canonical and disease and function pathways underlying the 144DG11 interactome were the lysosomal membranes (reference: GO:0005765 and KEGG pathway hsa04142). In parallel, STRING analysis (string-db.org) was used to visualize prominent nodes and enriched networks. For this first ranking of the compound interactome's specific proteins, we did not use the signal intensity of the peptides sequenced by MS because 1) the intrinsic properties of the technology cannot be based on protein quantitation (conversely to classic immunoprecipitation protocol for example), and 2) we do not use LC-MS/MS quantitative protocols (which would imply higher cost and longer time analysis). This unbiased analysis allowed Inoviem to classify potential relevant proteins and categorize them according to their involvement in specific pathways, or in relation with specific diseases. Following this bioinformatic selection, 8 proteins belonging to the autophagosomal-autolysosomal pathway were discovered (Fig. 5B). The discovery of this well-defined and enriched network demonstrates the overall success of the NPOT® experiment.

*Multi-parameter metabolic assessment.*

Metabolic and activity profiles of the mice were assessed by using the Promethion High-Definition Behavioral Phenotyping System (Sable Instruments, Inc., Las Vegas, NV, USA) as described previously *(46)*. Briefly, mice with free access to food and water were subjected to a standard 12 h light/12 h dark cycle, which consisted of a 48 h acclimation period followed by 24 h of sampling. Respiratory gases were measured by using the GA-3 gas analyzer (Sable Systems, Inc., Las Vegas, NV, USA) using a pull-mode, negative-pressure system. Air flow was measured and controlled by FR-8 (Sable Systems, Inc., Las Vegas, NV, USA), with a set flow rate of 2000 mL/min. Water vapor was continuously measured and its dilution effect on $O_2$ and $CO_2$ was mathematically compensated. Effective body mass was calculated by ANCOVA analysis as described previously *(47)*. Respiratory quotient (RQ) was calculated as the ratio of $VCO_2/VO_2$, and total energy expenditure (TEE) was calculated as $VO_2 \times (3.815 + 1.232 \times RQ)$, normalized to effective body mass, and expressed as kcal/h/$kg_{Eff.Mass}$. Fat oxidation (FO) and carbohydrate oxidation (CHO) were calculated as $FO = 1.69 \times VO_2 - 1.69 \times VCO_2$ and $CHO = 4.57 \times VCO_2 - 3.23 \times VO_2$ and expressed as g/d/$kg_{Eff.Mass}$. Activity and position were monitored simultaneously with the collection of the calorimetry data using XYZ beam arrays with a beam spacing of 0.25 cm. Food and water intakes were measured while calorimetric data was sampled.

*LAMP1 deglycosylation*

Mature human LAMP1 consists of a 354-amino acid (aa) luminal domain, a 23 aa transmembrane segment, and a 12 aa cytoplasmic tail. Its luminal domain is organized into two heavily N-glycosylated regions separated by a Ser/Pro-rich linker that carries a minor amount of O-linked glycosylation. Possible interaction between 144DG11 and a non-glycosylated form of LAMP1 was tested in order to establish whether the observed slow $k_{on}$ at pH 4.5 (calculated to be 1.31/millisecond, Fig. 5D) can be explained by diffusion interference of the heavy LAMP1 glycosylation. Deglycosylation was performed chemically by trifluoromethanesulfonic acid (TFMS) which completely removes all N- and O-linked glycans while preserving the protein structure. For the deglycosylation experiments, 25 μg of glycosylated LAMP1 luminal part was dialyzed, lyophilized, and then deglycosylated using manufacturer's recommendations (GlycoProfile™ IV Chemical Deglycosylation Kit, Sigma-Aldrich). RNase B (provided in the deglycosylation kit) was used as a positive control for a glycosylated protein. Finally, after short (24h) or long (72h) dialysis, the glycosylation status of LAMP1 and RNase B was tested by 15% SDS-PAGE mobility shift gel stained with QC colloidal Coomassie stain (#1610803, Bio-Rad). Our results (Fig. 5E, Upper panel) demonstrate a full deglycosylation of LAMP1 luminal part, with a unique band appearing after a long dialysis. RNAse B was also efficiently deglycosylated confirming the successful completion of the experiment. Glycosylated native LAMP1 protein, produced in mouse myeloma cells, migrated at around 110 kDa (Fig. 5E, lane 1), as expected from its multiple N- and O-glycosylation status. Upon TFMS treatment (Fig. 5E, lanes2 and 3), LAMP1 migrated at ≈ 40 kDa with no band at 110 kDa, indicating that 100% of the protein was efficiently deglycosylated. Long dialysis (Fig. 5E, lanes 3 and 6) enriched the deglycosylated bands. (B) A sensorgram showing the absence of interaction between deglycosylated LAMP1-Nter protein (degLAMP1-Nt) and 144DG11 (OKMW-XX1). degLAMP1-Nt was immobilized on the sensor chip CM5 using an alternative protocol without ethanolamine (Chip 2) and 144DG11 was diluted in HBS-EP pH 4 at the indicated concentrations, while the running buffer was at pH 5. The sensorgrams correspond to normalized signal (meaning subtracted of reference surface and buffer signal).

*Computational docking analysis*

LAMP1 sequence is divided into six segments (*18*): 1. residues M1-A28: signal sequence; 2. residues A29-R195: N-terminal domain; 3. residues P196-S216: linker between the domains; 4. residues S217-M382: C-terminal domain; 5. residues E383-V405: the transmembrane segment; residues G406-I417: cytoplasmic domain. We have analyzed only the N- and the C-terminal domains since: 1. The signal sequence, transmembrane segment and the cytoplasmic domain are assumed to be irrelevant for the binding of small molecules; 2. The linker between the domains is unstructured and heavily glycosylated (7 out of 20 residues) and thus too complicated to model. We have not considered glycosylation in the N- and the C-terminals. The C- and N-Terminal domains were modeled based on the known crystal structure of mouse LAMP1 C-terminal domain (PDB ID 5gv0) which is structurally highly similar to the N-terminal domain (*18*). The MODELLER software tool *(48)* was used for homology modeling, producing 5 optional models for each domain. The obtained 10 models (as well as 5gv0 itself) were prepared in pH 5 by the "protein preparation wizard" as implemented in Schrodinger 2020-2. Possible binding sites were identified by three different computational tools: SiteMap *(49)*, FtSite *(50)* and fPocket.*(51)*. Overall, 130 optional sites were identified in 11 LAMP1 3D structures. Docking computations were performed for each of the putative binding sites: 418 out of a large and diverse database of ~30 million molecules were chosen as decoys according to 144DG11 applicability domain (Lipinski rules properties). The decoys library was narrowed down to 233 based on chemical similarity (Tanimoto coefficient >=0.7). Docking computations for 144DG11 in a set of molecules composed of 144DG11 and 233 decoys (prepared in pH 5) were performed for every putative binding site in every model (overall 130 sites). The computations were performed using the Glide algorithm *(52)*, as implemented in Schrodinger 2020-2. According to the docking results analysis (Data file S2), in 18 out of 130 sites 144DG11 was ranked at the top 10% (rankings 1-24, Data file S2). Analyzing the results, we realized that site 1 of SiteMap, site 3 of fPocket, and site 2 of FtSite refer to the same pocket (residues F50-D55, N62, L67, F118, Y120-L122, T125, L127-S133, N164-V166).

We examined the differences between the binding modes of 144DG11 to the common site as defined by each software tool (Fig. 5F) and came to the conclusion that 144DG11 tends to bind in a similar binding mode regardless of the software by which the site was configured (two out of three binding modes (SiteMap and fPocket) were identical while in FtSite, part of the molecule went through a rotation relative to the other two, but overall, the poses were highly similar).

To ensure 144DG11 specificity to the identified binding site, we repeated the analysis presented above for all 233 decoys. Only in 13 out of 233 molecules, we observed the same results as for 144DG11 - *i.e.,* the molecules were successfully docked to pockets predicted by all three tools. Moreover, the pocket identified for 144DG11 (Fig. 5E) matched 4 molecules out of 13). From these results, we shall deduce that the molecule 144DG11 selectively binds LAMP1 at the identified binding site.

In summary, we have computationally identified a possible binding site for 144DG11 in the N-terminal domain of LAMP1 and predicted with high certainty that this result is specific for 144DG11 since the probabilities to obtain similar results for decoy molecules are low.

*Transmission electron microscopy (TEM)*

Liver tissue was minced and fixed in a solution containing 2% paraformaldehyde, 2.5 % glutaraldehyde (EM grade) in 0.1M sodium cacodylate buffer pH 7.3 for 2 hours at RT, followed by 24 h at 4°C. Tissue was then washed 4 times with sodium cacodylate and postfixed for 1 h with 1% osmium tetroxide and 1.5% potassium ferricyanide in sodium cacodylate. Then sample was washed 4 times with the same buffer and dehydrated with graded series of ethanol solutions (30, 50, 70, 80, 90, 95 %) for 10 minutes each and then 100% ethanol 3 times for 20 minutes each. Subsequently, samples were treated with 2 changes of propylene oxide. Samples were then infiltrated with series of epoxy resin (25, 50, 75, 100% - 24 h in each) and polymerized in the oven at 60°C for 48 hours. The blocks were sectioned by an ultramicrotome (Ultracut E, Riechert-Jung) and obtained sections of 80 nm were stained with uranyl acetate and lead citrate. Sections were observed by Jeol JEM 1400 Plus Transmission Electron Microscope and images were taken using Gatan Orius CCD camera.

*Proteomics (Fig. 7)*

*Sample preparation for MS analysis.* Cell lysates in RIPA buffer containing protease inhibitors were clarified by centrifugation and 40 µg of protein was used for protein precipitation by the chloroform/methanol method (*53*). The precipitated proteins were solubilized in 100 µl of 8M urea, 10 mM DTT, 25 mM Tris-HCl pH 8.0 and incubated for 30 min at 22°C. Iodoacetamide (55 mM) was added and samples were incubated for 30 min (22°C, in the dark), followed by addition of DTT (10 mM). Fifty µl of the samples was transferred into a new tube, diluted by the addition of 7 volumes of 25 mM Tris-HCl pH 8.0 and sequencing-grade modified Trypsin (Promega Corp., Madison, WI) was added (0.35 µg/ sample) followed by incubation overnight at 37°C with gentle agitation. The samples were acidified by addition of 0.2% formic acid and desalted on C18 home-made Stage tips. Peptide concentration was determined by Absorbance at 280 nm and 0.75 µg of peptides were injected into the mass spectrometer.

*nanoLC-MS/MS analysis*. MS analysis was performed using a Q Exactive-HF mass spectrometer (Thermo Fisher Scientific, Waltham, MA USA) coupled on-line to a nanoflow UHPLC instrument, Ultimate 3000 Dionex (Thermo Fisher Scientific, Waltham, MA USA). Peptides dissolved in 0.1% formic acid were separated without a trap column over a 120 min acetonitrile gradient run at a flow rate of 0.3 µl/min on a reverse phase 25-cm-long C18 column (75 µm ID, 2 µm, 100Å, Thermo PepMapRSLC). The instrument settings were as described in (*54*). Survey scans (300–1,650 m/z, target value 3E6 charges, maximum ion injection time 20 ms) were acquired and followed by higher energy collisional dissociation (HCD)-based fragmentation (normalized collision energy 27). A resolution of 60,000 was used for survey scans and up to 15 dynamically chosen most abundant precursor ions, with "peptide preferable" profile were fragmented (isolation window 1.6 m/z). The MS/MS scans were acquired at a resolution of 15,000 (target value 1E5 charges, maximum ion injection times 25 ms). Dynamic exclusion was 20 sec. Data were acquired using Xcalibur software (Thermo Scientific). To avoid a carryover, the column was washed with 80% acetonitrile, 0.1% formic acid for 25 min between samples.

*MS data analysis.* Mass spectra data were processed using the MaxQuant computational platform, version 1.6.14.0. Peak lists were searched against the Uniprot human FASTA sequence database from May 19, 2020 containing 49,974 entries. The search included cysteine carbamidomethylation as a fixed modification, N-terminal acetylation and oxidation of methionine as variable modifications and allowed up to two miscleavages. The match-between-runs option was used. Peptides with a length of at least seven amino-acids were considered and the required FDR was set to 1% at the peptide and protein level. Relative protein quantification in MaxQuant was performed using the label-free quantification (LFQ) algorithm (*55*). Statistical analysis (n=4-7) was performed using the Perseus statistical package (*56*). Only those proteins for which at least 3 valid LFQ values were obtained in at least one sample group were accepted for statistical analysis by t-test ($p < 0.05$).

**Supplementary figures**

**Fig. S1.** *In silico* ADMET (Absorption, Distribution, Metabolism, and Excretion Toxicity)-compatible, polyglucosan lowering compounds.

Table shows analysis of three different ADMET algorithms. On the left are ordinal numbers of the compounds according to the hits discovered in (*13*). A range of numbers refers to enantiomers. The second column from left is the ChemBridge catalog number of the compound and the Heat map demonstrates the number of violations predicted by each algorithm

| | Candidate | | # of problems per program | | | Decission |
|---|---|---|---|---|---|---|
| | | | QikProp | SwissADME | AdmetSAR | |
| | 01-02 | 86282818 | 4 | 3 | 5 | |
| | 03-06 | 15607447 | 4 | 4 | 6 | |
| | 07 | 42992072 | 1 | 2 | 5 | |
| | 08-09 | 25760823 | 4 | 1 | 4 | |
| | 10-11 | 82320451 | 2 | 0 | 5 | |
| | 12 | 42459198 | 1 | 2 | 4 | |
| | 13-14 | 37867671 | 1 | 2 | 5 | |
| 144-DG-11 → | 15-16 [A] | 27686904 | 2 | 1 | 3 | Preferred |
| | 17-18 | 17057751 | 2 | 2 | 5 | |
| | 19 | 38058095 | 2 | 0 | 2 | Preferred |
| | 20-23 | 36585388 | 4 | 2 | 6 | |
| | 24 [B] | 88095528 | 3 | 1 | 5 | |
| | 25 | 57540036 | 1 | 4 | 4 | |
| | 26 | 54056378 | 4 | 2 | 4 | |
| | 27-28 | 83101459 | 2 | 2 | 3 | Preferred |
| | 29 | 76195865 | 1 | 0 | 3 | Preferred |
| | 30 | 34834825 | 3 | 1 | 4 | |
| | 31-32 | 68349003 | 3 | 1 | 4 | |
| | 33-36 | 78653061 | 3 | 1 | 6 | |

**Fig. S2.** An ADMET-incompatible compound (88095528 in Fig. S1) causing wounds in Gbe$^{ys/ys}$ mice.

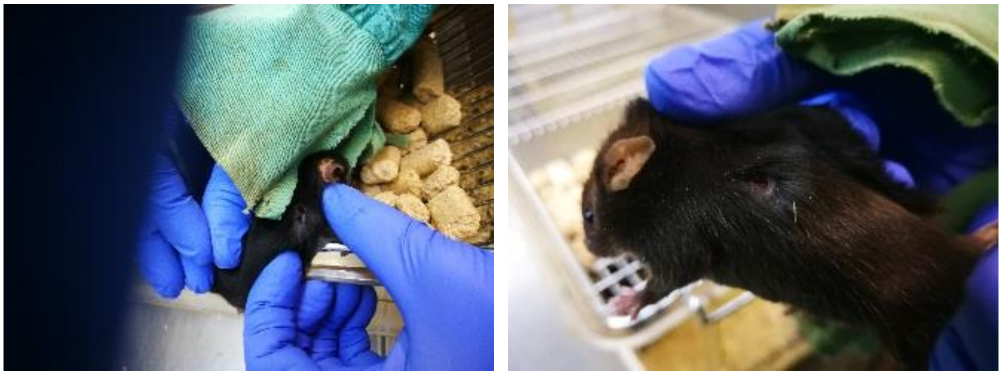

**Fig. S3.** Body weights of wild type C57Bl6J mice treated with 144DG11 for 3 months.

Mice were injected twice a week with 150 uL of 144DG11 at 250 mg/kg in 5% DMSO (red), or an equal volume of 5% DMSO (V, vehicle) control (black). Injections were intravenous for the first month and then subcutaneous for the following 2 months. No significant change between the two treatments is observed.

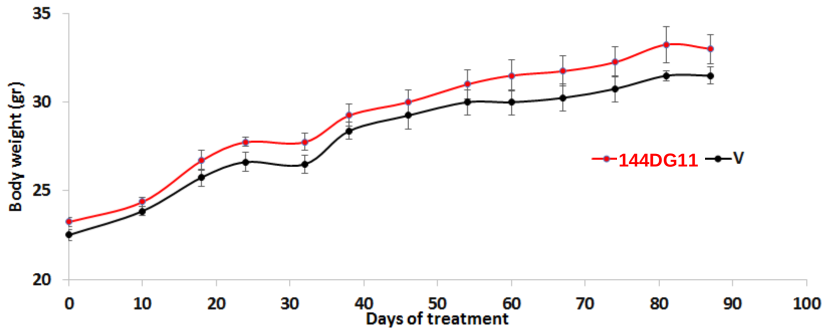

**Fig. S4.** Histology of 144DG11 tissues compared to vehicle control.

Brain, liver, skeletal muscle and heart slices of wild type C57Bl6J mice treated for 3 months with 144DG11 as in Fig. S3. The slices were stained by H&E staining in order to visualize lesions. No lesions were apparent in either treatment. Scale bars, 500 µm (brain), 100 µm (liver), 200 µm (muscle), 100 µm (heart).

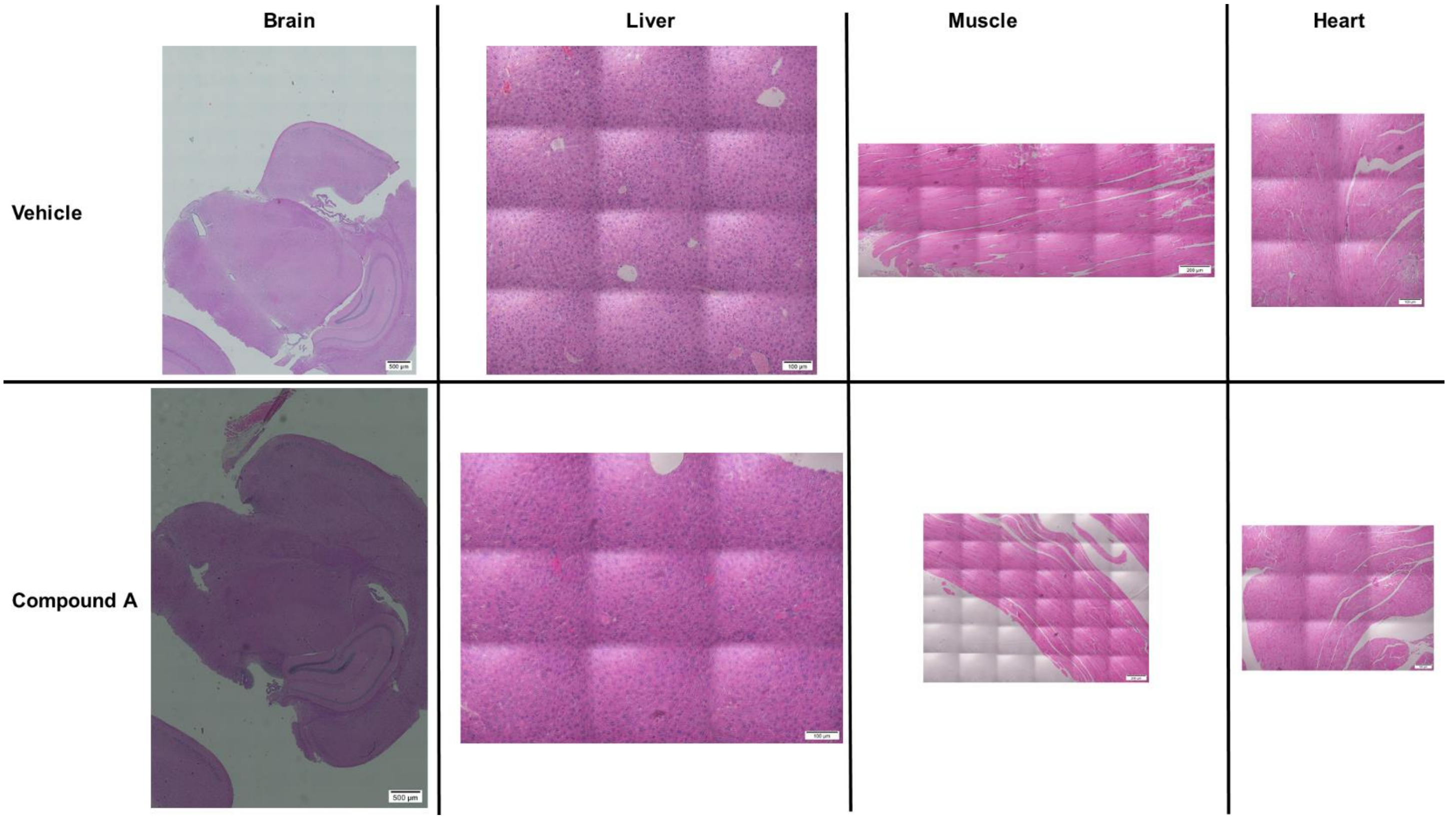

**Fig. S5.** Therapeutic strategies for APBD.

Therapeutic approaches for APBD are based on reduction of the GYS/GBE activity ratio or on direct PG and glycogen degradation. Blue hexagon, glucose; black hexagon, glucose-1-phosphate.

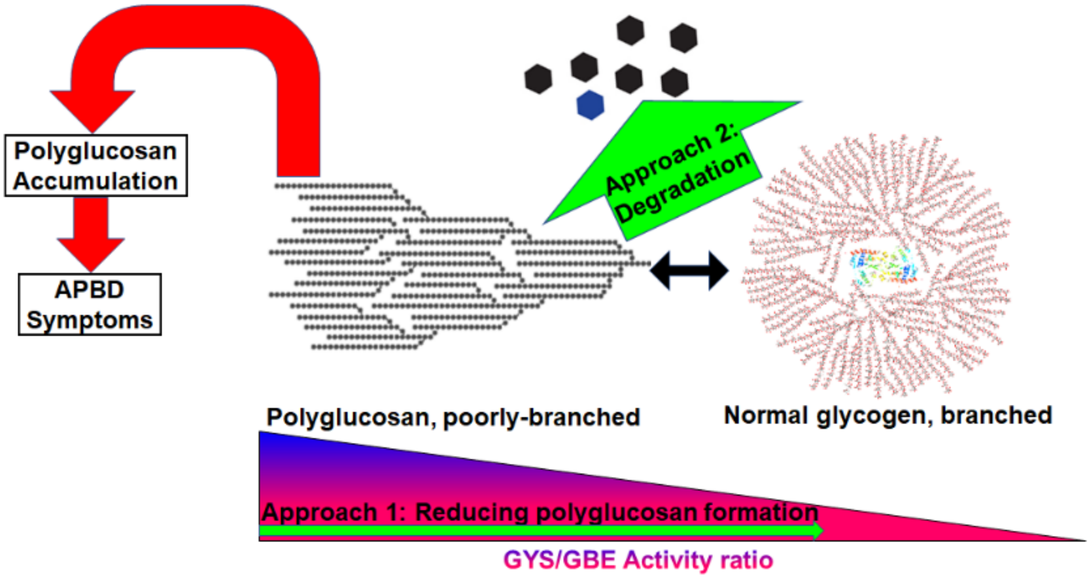

**Fig. S6.** 144DG11 at the LAMP1:LAMP1 interface.

To evaluate the probability that 144DG11 can interfere with LAMP1:LAMP1 interactions by binding to the predicted binding site at the LAMP1 N-terminus, we performed LAMP1 N-terminus:LAMP1 N-terminus protein:protein docking computations **(A)**. According to the three highest ranked solutions (the top ranked result is shown in **(A)**), 144DG11 putative binding site is located in the LAMP1:LAMP1 interface. The possibility that 144DG11 also inhibits LAMP1:LAMP2 interactions requires additional computations. **(A)** Predicted binding site for 144DG11 in LAMP1's N-terminal domain. Top ranked solution obtained by PATCHDOCK and FireDock servers. 144DG11, represented by black sticks, was docked to the putative binding site. LAMP1 N-terminal chains are represented in green and cyan. **(B)** Schematic of the lysosomal membrane (LM), LAMP1, LAMP2 and the potential inhibitor 144DG11. **?**, a possible ancillary membrane protein mediating LAMP1 interaction.

A

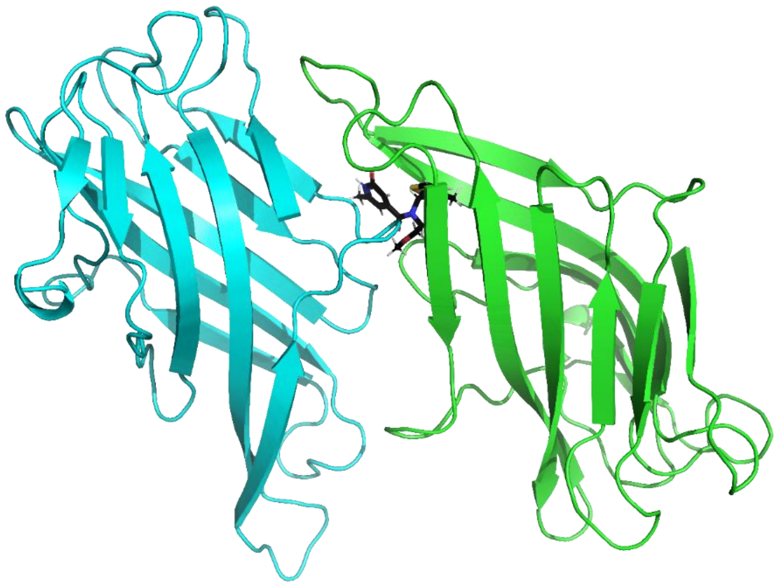

B

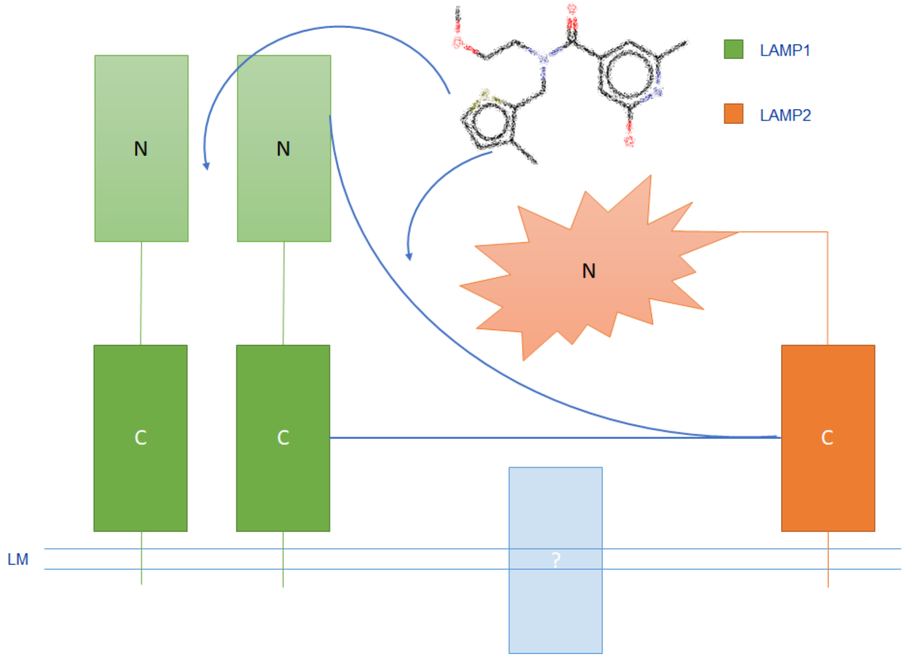

**Fig. S7.** Heteroassemblies formed around 144DG11.

Photos of the heteroassemblies (red circles) obtained by NPOT® on APBD-patient fibroblasts **(A)**, or HC fibroblasts **(B)** in the presence of compounds 144DG11 (OKMW-XX1) and OKMW-XXC (negative control) at $10^{-6}$ M. Each experiment was done in triplicate. Technical negative controls are obtained without the addition of any compound. Each picture represents a well of a 96-well plate.

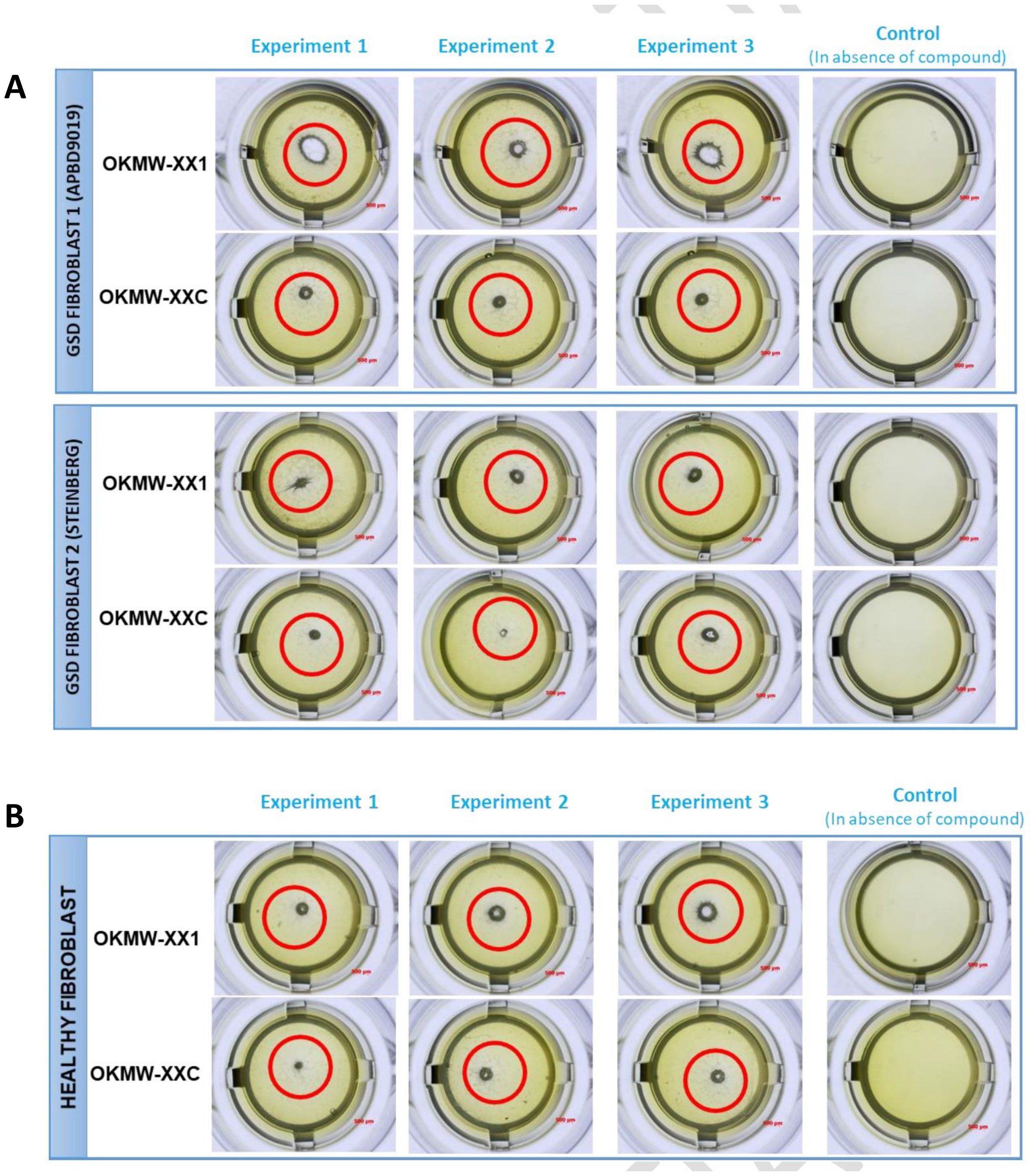

**Fig. S8.** Bioenergetic parameters of 144DG11 treated cells.

**(A)** Basal respiration in the indicated groups calculated as the mean OCR from the initiation of the experiment until first injection of oligomycin. 144DG11 led to a significant increase in basal respiration in both healthy control ($p<0.007$) and APBD patient ($p<0.02$) cells. **(B)** Maximal respiration, defined as the difference between OCR values after FCCP and rotenone/antimycin supplementations, is increased by 144DG11 in both HC ($p<0.04$) and APBD patient ($p<0.03$) cells. **(C)** ATP production, defined as the difference in OCR values between basal and post oligomycin levels, is increased by 144DG11 only in APBD patient cells ($p<0.006$). **(D)** Spare respiratory capacity, defined as the difference between maximal (post FCCP) OCR and basal OCR values, was slightly (not significantly) increased by 144DG11 in both HC ($p<0.1$) and APBD patient ($p<0.13$) cells. **(E)** Coupling efficiency, defined as the quotient (OCR following Oligomycin)/(Basal OCR), was increased by 144DG11 in HC ($p<0.001$) and APBD patient ($p<0.02$) cells. All analyses are based on mean values. Statistical analysis was done by One-Way ANOVA with Dunnett post-hoc tests.

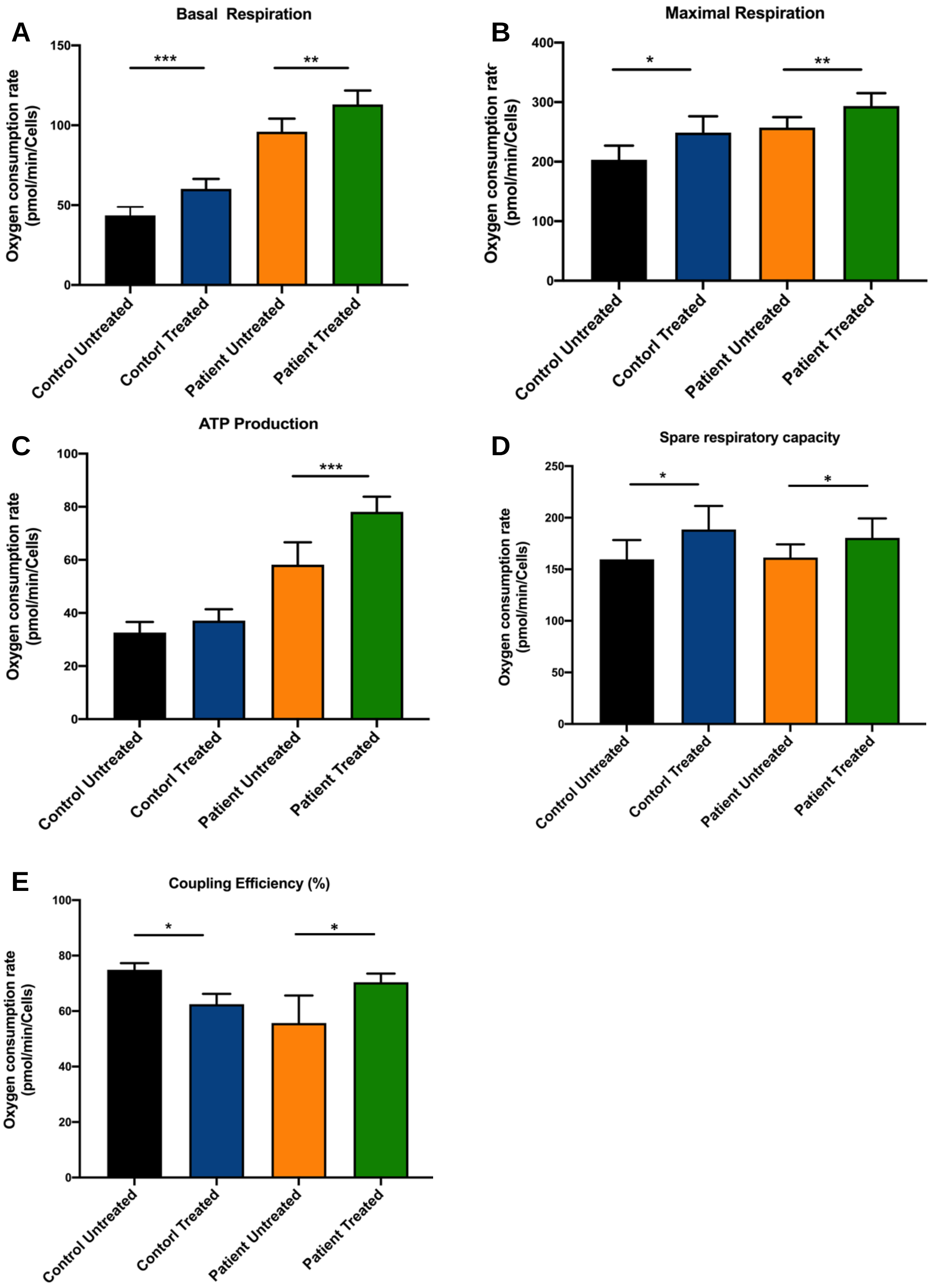

**Fig. S9.** Endocytosis – a KEGG-Pathway enriched in APBD patient fibroblasts.

Proteins up-regulated in APBD fibroblasts, as compared to HC (Figure 7E) were analyzed by the KEGG Pathway annotation tool in DAVID. The endocytosis pathway was found to be enriched in APBD fibroblasts ($p<1.10^{-6}$, $FDR<2.1x10^{-4}$). Stars denote proteins significantly modulated by the APBD diseased state.

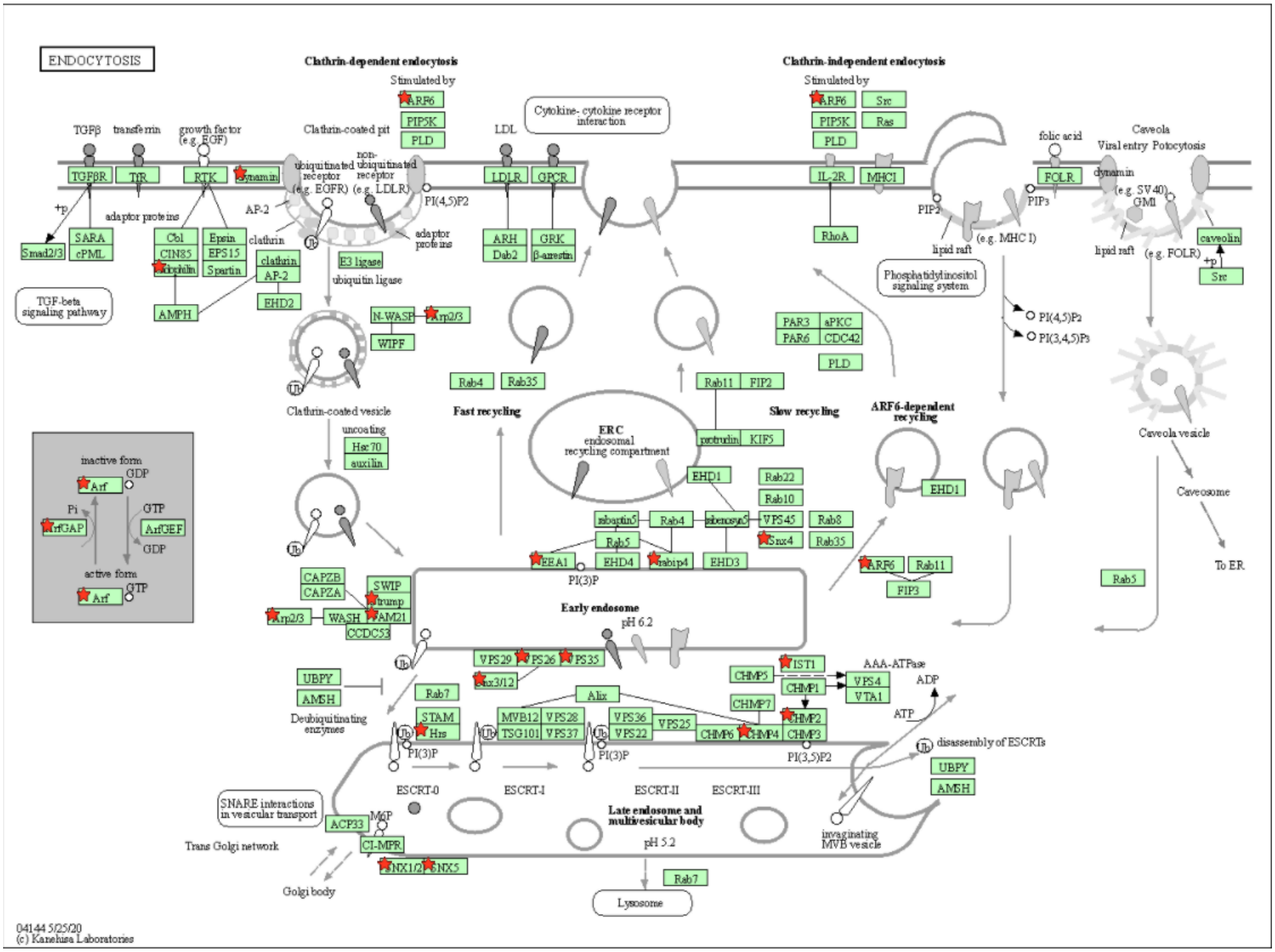

**Fig. S10.** Oxidative Phosphorylation – a KEGG Pathway depleted in APBD fibroblasts.

Proteins down-regulated in APBD fibroblasts, as compared to HC (Figure 7E) were analyzed by the KEGG Pathway annotation tool in DAVID. The oxidative phosphorylation pathway was found to be depleted in APBD fibroblasts ($p<1.9\times10^{-8}$, $FDR<1.5\times10^{-6}$). Stars denote proteins significantly down-modulated by the APBD diseased state.

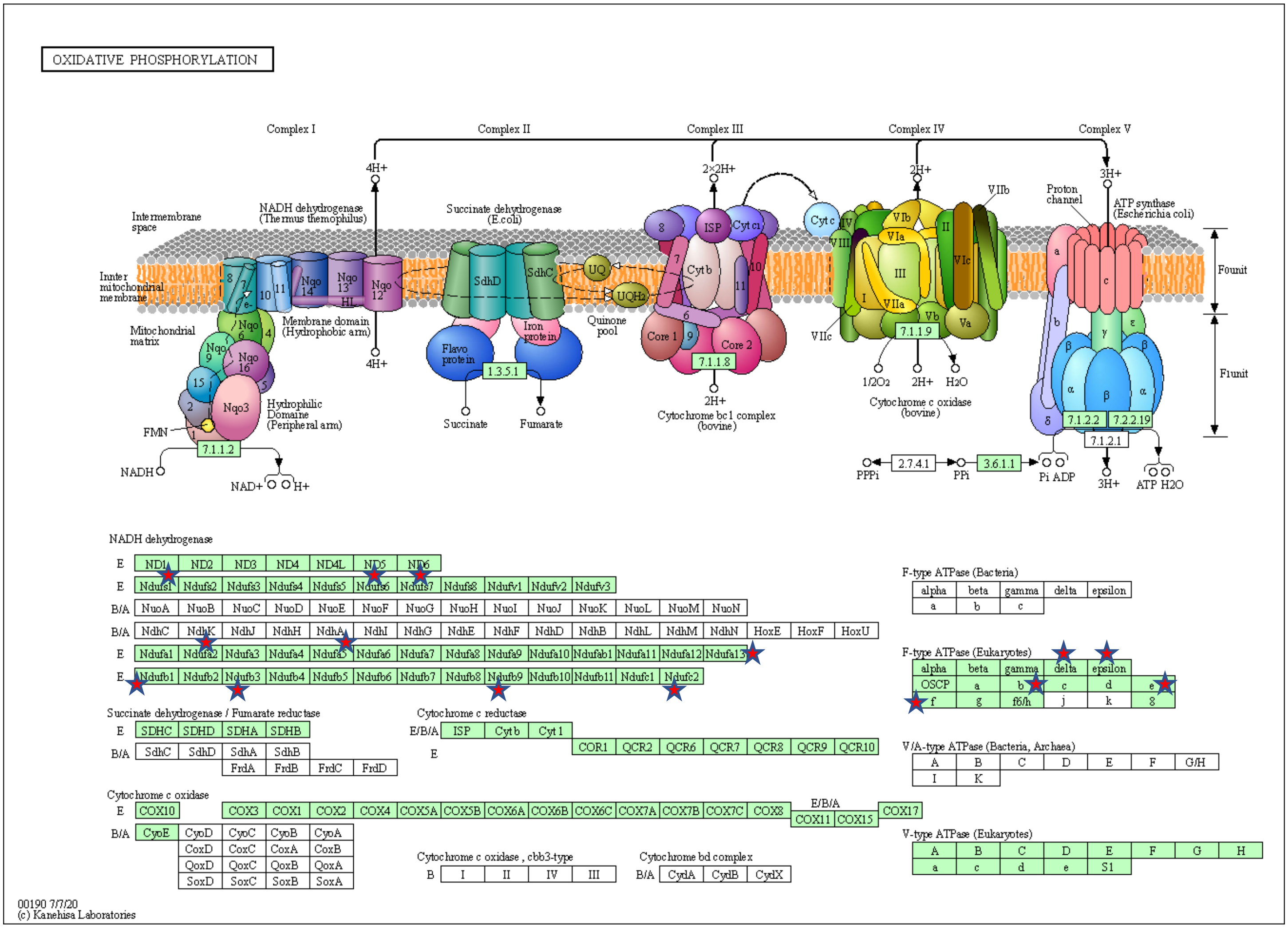

**Table S1.** Irwin test results of 144DG11.

| | Vehicle | 50 mg/kg 1h | 250 mg/kg 1h | 50 mg/kg 24h | 250 mg/kg 24h |
|---|---|---|---|---|---|
| Coat color | Black | Black | Black | Black | Black |
| Presence of whiskers | 3 | 3 | 3 | 3 | 3 |
| Appearance of fur | 2 | 2 | 2 | 2 | 2 |
| Piloerection | 0 | 0 | 0 | 0 | 0 |
| Patches of missing fur on face | 0 | 0 | 0 | 0 | 0 |
| Patches of missing fur on body | 0 | 0 | 0 | 0 | 0 |
| Wounds | 0 | 0 | 0 | 0 | 0 |
| Transfer behavior | 5 | 5 | 5 | 5 | 5 |
| Body position | 3.5 | 3.5 | 3 | 3.5 | 3 |
| Tremor | 0 | 0 | 0 | 0 | 0 |
| Gait | 0 | 0 | 0 | 0 | 0 |
| Pelvic elevation | 2 | 2 | 2 | 2 | 2 |
| Tail elevation | 1 | 1 | 1 | 1 | 1 |
| Touch escape | 2 | 2 | 2 | 2 | 2 |
| Positional passivity | 0 | 0 | 0 | 0 | 0 |
| Trunk curl | 0 | 0 | 0 | 0 | 0 |
| Righting reflex | 0 | 0 | 0 | 0 | 0 |
| Salivation | 0 | 0 | 0 | 0 | 0 |
| Extension reflex | 2 | 2 | 2 | 2 | 2 |

**Irwin test scores:** *Presence of whiskers* – 0 = None, 1 = A few, 2 = Most, but not a full set, 3 = A full set; *Appearance of fur* – 0 = Ungroomed and disheveled, 1 = Somewhat disheveled, 2 = Well-groomed (normal); *Piloerection* – 0 = None, 1 = Most hairs standing on end; *Patches of missing fur on face* – 0 = None, 1 = Some, 2 = Extensive; *Patches of missing fur on body* - 0 = None, 1 = Some, 2 = Extensive; *Wounds* – 0 = None, 1 = Signs of previous wounding, 2 = Slight wounds present, 3 = Moderate wounds present, 4 = Extensive wounds present; *Transfer behavior* – 0 = Coma, 1 = Prolonged freeze (>10 sec) then slight movement, 3 = Brief freeze (a few seconds) then active movement, 4 = Momentary freeze then swift movement, 5 = No freeze immediate movement, 6 = Extremely excited (manic); *Body position* – 0 = Completely flat (on stomach), 1 = Lying on side, 2 = Lying on back, 3 = Sitting or standing, 4 = Rearing on hind legs, 5 = Repeated vertical leaping; *Tremor* – 0 = None, 1 = Mild, 2 = Marked; *Gait* – 0 = Normal, 1 = Fluid but abnormal, 2 = Limited movement only, 3 = Incapacity; *Pelvic elevation* – 0 = Markedly flattened, 1 = Barely touches, 2 = Normal (3 mm elevation), 3 = Elevated (more than 3 mm elevation); *Tail elevation* – 0 = Dragging, 1 = Horizontally extended, 2 = Elevated (Straub tail); *Touch escape* – 0 = No response, 1 = Mild (escape response to firm stroke), 2 = Moderate (rapid response to light stroke), 3 = Vigorous (escape response to approach); *Positional passivity* – 0 = Struggles when restrained by tail, 1 = Struggles when restrained by neck (finger grip, not scruffed), 2 = Struggles when held supine (on back), 3 = Struggles when restrained by hind legs, 4 = Does not struggle; *Trunk curl* – 0 = Absent, 1 = Present; *Righting reflex* – 0 = No impairment, 1-10 = Number of seconds required to right; *Salivation* – 0 = None, 1 = Slight margin of sub-maxillary area, 2 = Wet zone entire sub-maxillary area; *Extension reflex* – 0 = Severe defect in hind limb extension reflex, 1 = Mild defect in hind limb reflex, 2 = Normal.

**Table S2.** Decoy molecules docked to pockets predicted by 3 computational tools.

Thirteen decoys were successfully docked to several pockets, predicted by all three tools. SiteMap, FtSite, and fPocket.

| Model, Site ranking | Compounds (InchiKeys) | Number of compounds |
|---|---|---|
| **5gv0, site112** | ISNBQFVIRRXMNN-UHFFFAOYSA-N | 1 |
| **HN.B99990001, site113** | SDIOOFTYDJNAOO-UHFFFAOYSA-N | 1 |
| **HN.B99990004, site123** | **144DG11**<br>AFRQZZBRQMVSOV-UHFFFAOYSA-N*<br>OYOBHSNWLQNJLP-UHFFFAOYSA-N<br>ULMGNNFJGOAZQX-UHFFFAOYSA-N<br>VODWQUHDWRIEOI-UHFFFAOYSA-N | 5 |
| **HN.B99990005, site121** | LHKJBSUPOYJYCL-UHFFFAOYSA-N | 1 |
| **HN.B99990005, site233** | UIVJRUWFXCGSSM-UHFFFAOYSA-N<br>XDGVKHPNDVKOPJ-UHFFFAOYSA-N | 2 |
| **HC.B99990001, site223** | OHHMZSKNHGGUAX-UHFFFAOYSA-N | 1 |
| **HC.B99990002, site131** | CKHGBJKQAHAIDZ-UHFFFAOYSA-N<br>GLXDFBGZVSRFGI-UHFFFAOYSA-N<br>WTTMQZWWJFRCSI-UHFFFAOYSA-N | 3 |
| **HC.B99990003, site233** | AFRQZZBRQMVSOV-UHFFFAOYSA-N* | 1 |

* Note: this molecule binds to two different binding sites, indicating is probably promiscuous.

**Data file S1**. Proteomics raw data

**Data file S2.** LAMP1 docking sites to 144DG11

**Movie S1.** Motor effects of 144DG11 in Gbe$^{ys/ys}$ mice.
